# In the non-canonical NF-κB pathway, binding of the initiating kinase NIK to the pathway’s target, p100, serves a nonenzymatic “licensing” role

**DOI:** 10.64898/2026.11.16.766979

**Authors:** Xun Liang, David Scheerer, Konstantin Bogdanov, Mike Gardarsky, Chaitanyakumar Amballa, Tianzhen Wang, David Morgenstern, Moshe Goldsmith, Dana Reichmann, Gilad Haran, David Wallach

## Abstract

The non-canonical NF-_κ_B pathway plays pivotal roles in adaptive immunity. It is initiated by NIK-mediated phosphorylation of IKK1, which phosphorylates p100 and promotes processing of newly synthesized p100. However, IKK1 can also be activated by other kinases to mediate distinct functions. Why IKK1-induced p100 processing occurs efficiently downstream of NIK but not of other kinases is unknown. Here, we show that whereas pre-existing p100 can be phosphorylated by IKK1 without NIK, phosphorylation of newly synthesized p100 critically depends on additional non-enzymatic functions of NIK’s C-terminal portion. NIK binds IKK1 and restrains its kinase activity, while its binding to p100 enhances p100 phosphorylation. Single-molecule FRET spectroscopy and XL-MS revealed that the C-terminal region of p100 adopts a folded-back conformation and undergoes structural rearrangement upon NIK binding, likely accounting for this enhancement. Thus, NIK not only initiates the pathway but also dictates its specificity through binding to its downstream substrate.

## INTRODUCTION

Phosphorylation of serine residues S866 and S870 at the C terminus of the NF-_κ_B precursor protein NF-_κ_B2 (p100) by the protein kinase IKK1 plays a pivotal role in the regulation of adaptive immunity. It directs the processing of p100 into the NF-_κ_B protein p52, which, in association with RelB or other NF-_κ_B proteins, activates numerous immune regulatory genes. This phosphorylation is dictated by the so-called ‘non-canonical NF-_κ_B signaling pathway’. In this pathway, which is primarily initiated by certain members of the TNF ligand family, receptor stimulation increases the cellular level of a MAP3K called NIK, which in turn activates IKK1 by phosphorylating it ^1–4^.

However, IKK1 can also be activated by other mechanisms to perform functions distinct from those mediated by the non-canonical NF-_κ_B pathway. In response to signaling pathways that do not involve NIK, IKK1 acts either alone or in complex with the related kinase IKK2 to regulate targets other than p100, thereby controlling a variety of biological processes distinct from those governed by NIK (e.g.,^5–9^). Some of those important functions of IKK1 are exerted independently of its kinase activity (e.g.,^10^).

Why only IKK1 molecules that were activated by NIK dictate processing of p100 to p52 remains poorly understood.

It has been previously shown that only newly synthesized p100 molecules serve as potential precursors of p52 ^11^. p100 molecules that have accumulated in the cell for some time (pre-existing p100) serve another important functional role. They assemble with other NF-_κ_B proteins into high-molecular-weight _κ_Bsome complexes that sequester NF-_κ_B proteins and inhibit their activity ^12–14^. Exploring the nature of the specificity in the effect of IKK1 on p100, we have found in the present study that S866 and S870 in pre-existing p100 can also be phosphorylated by IKK1 activated independently of NIK. However, IKK1-mediated phosphorylation of p100 is substantially more efficient in the presence of NIK than in its absence. Importantly, phosphorylation of newly synthesized p100 was found to strictly depend on NIK.

Exploring the mechanism by which NIK dictates this specificity, we found that, in addition to activating IKK1 enzymatically, NIK promotes IKK1-mediated phosphorylation of p100 through an additional, non-enzymatic, mechanism. This effect does not result from further enhancement of the kinase activity of phosphorylated IKK1. In fact, we found that persistent association of NIK with phosphorylated IKK1 restricts the kinase activity of the latter. Rather, NIK acts directly on p100, inducing a conformational rearrangement that likely accounts for NIK-mediated facilitation of p100 phosphorylation.

Thus, NIK not only activates IKK1 by phosphorylating it, but also functions as a non-enzymatic specificity factor that licenses IKK1-mediated phosphorylation of newly synthesized p100.

## RESULTS

### NIK non-enzymatically promotes IKK1-mediated phosphorylation of p100 at S866 and S870

When the protein kinase IKK1 is activated in response to ligands that upregulate the expression of the MAP3K NIK (in the so-called ‘non-canonical NF-κB pathway’), it phosphorylates p100 at S866 and S870, thereby recruiting βTrCP and initiating p100 processing. However, IKK1 can also be activated by kinases other than NIK (e.g., in the canonical NF-κB pathway), where it serves other functions. This raises the question of how upregulation of NIK expression specifically directs IKK1 toward p100 phosphorylation and processing.

To address this question, we compared the effect of NIK on IKK1 with that of TAK1, a MAP3K that activates IKK1, as well as IKK2, which functions in the canonical NF-κB pathway ^15^. As shown in Fig. 1A, expression of a fusion protein consisting of the kinase domain of TAK1 fused to the minimal TAK1-activating domain of the adaptor protein TAB1 (TAK1-TAB1) ^16^ induced phosphorylation of p100 at S866 and S870, as did expression of NIK. This response, like that induced by NIK, was completely dependent on IKK1 (Fig. S1A). However, relative to the extent of IKK1 phosphorylation, TAK1 induced much less phosphorylation of p100 than NIK (Fig. 1A and B). Moreover, whereas NIK-induced phosphorylation promoted processing of p100 to p52, TAK1–TAB1-induced phosphorylation did not, even when TAK1-TAB1 expression was sufficient to induce levels of S866/S870 phosphorylation comparable to those induced by NIK (Fig. 1C). IL-1β, a pro-inflammatory cytokine that activates TAK1, has likewise been reported to induce phosphorylation of p100 at S866/S870 without promoting its processing ^17^. The same outcome was also observed following stimulation by the pro-inflammatory cytokine TNF (Figs. 1D and S1B).

**Figure 1.**
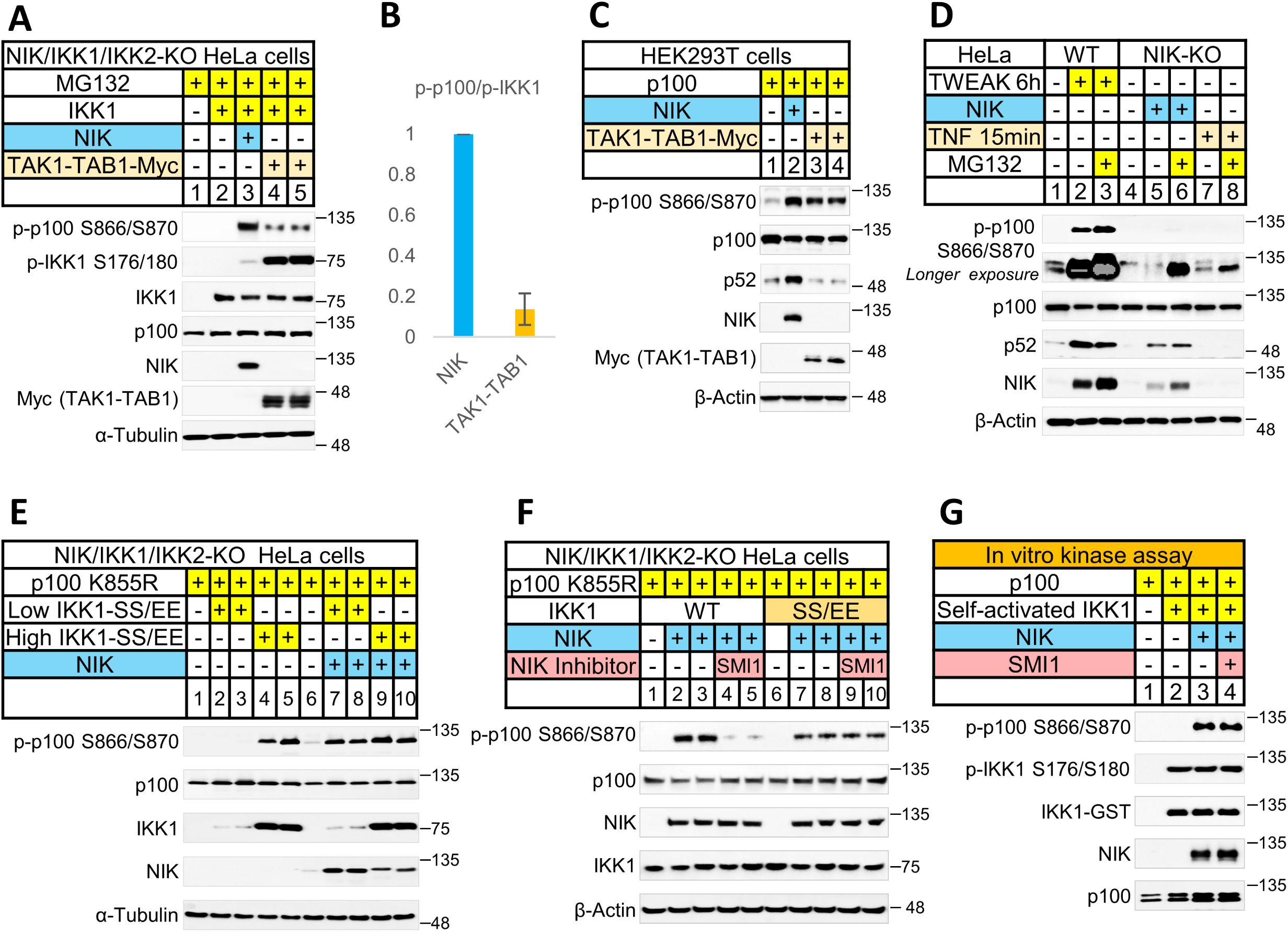
NIK non-enzymatically enhances IKK1-mediated phosphorylation of p100 at S866 and S870. A. Immunoblot analysis of IKK1 and p100 phosphorylation in NIK/IKK1/IKK2-knockout HeLa cells co-transfected with a plasmid encoding IKK1 together with either NIK or TAK1-TAB1. MG132 (10 μM) was added 1 h before analysis, as indicated. Lanes 4 and 5 are independently prepared replicate samples from separate wells within the same experiment. B. Bar chart showing the relative ratio of phosphorylated p100 to phosphorylated IKK1 in the NIK- and TAK1–TAB1-expressing samples shown in (A), with the NIK-expressing sample set to 1. The TAK1–TAB1 bar represents the mean ± SD of the two independently prepared replicate samples. C. Immunoblot analysis of p100 phosphorylation and processing in HEK293T cells co-transfected with a plasmid encoding p100 together with a plasmid encoding either NIK or TAK1-TAB1. D. Immunoblot analysis of p100 phosphorylation and processing in wild-type HeLa cells following stimulation with TWEAK (100 ng/ml), or in NIK-knockout HeLa cells following stimulation with TNF (100 ng/ml) or ectopic expression of NIK. MG132 (10 μM) was added 1 h before analysis, as indicated. E. Immunoblot analysis of p100 phosphorylation in NIK/IKK1/IKK2-knockout HeLa cells co-transfected with plasmids encoding p100-K855R and IKK1-SS/EE, in the presence or absence of a plasmid encoding NIK. Low- and high-level expression of IKK1-SS/EE were achieved using pBABE and pLV expression vectors, respectively. F. Immunoblot analysis of p100 phosphorylation in NIK/IKK1/IKK2-knockout HeLa cells co-transfected with plasmids encoding p100-K855R and either wild-type IKK1 or IKK1-SS/EE. Cells were treated with SMI1 (1 μM) for 2 h before analysis. G. In vitro phosphorylation of recombinant p100 by self-activated IKK1 in the presence or absence of recombinant NIK (see Methods). SMI1 (200 nM) was added where indicated.

To explore the mechanism by which NIK enhances the effect of activated IKK1 on p100, we assessed the phosphorylation of S866/S870 in p100 by the constitutively active IKK1-S176E/S180E mutant (IKK1-SS/EE). As shown in Figs. 1E and S1C, high-level expression of IKK1-SS/EE in cells lacking endogenous IKK1 and NIK was sufficient to induce phosphorylation of S866/S870 in p100. However, when expressed at low levels, IKK1-SS/EE phosphorylated S866/S870 in p100 only in cells that co-expressed NIK.

Phosphorylation of p100 at S866/S870 induced by IKK1-SS/EE alone failed to promote processing of p100 to p52 (Fig. S1D), as observed in TNF-treated cells and in cells expressing TAK1-TAB1.

To determine whether NIK enhances the function of activated IKK1 through its kinase function, we treated the cells with SMI1, an inhibitor of NIK kinase activity. As shown in Fig. 1F, SMI1 strongly inhibited p100 phosphorylation in cells expressing wild-type IKK1 and NIK but had no effect on the NIK-dependent enhancement of p100 phosphorylation in cells expressing low levels of IKK1-SS/EE. Thus, this second function of NIK appears to be independent of its kinase activity.

To determine whether NIK directly enhances phosphorylation of p100 by activated IKK1, we reconstituted the reaction in vitro using recombinant proteins (Fig. 1G). Activated recombinant IKK1 efficiently phosphorylated recombinant p100 only when recombinant NIK was included in the reaction. Consistent with the cell-based experiments, SMI1 had no effect on the ability of NIK to enhance p100 phosphorylation in vitro.

Together, these findings demonstrate that NIK promotes p100 phosphorylation by two distinct mechanisms. In addition to activating IKK1 enzymatically, NIK exerts a second, non-enzymatic activity that enhances phosphorylation of p100 by activated IKK1 and enables this phosphorylation to drive p100 processing.

### IKK1-mediated phosphorylation of newly synthesized p100, but not of pre-existing p100 molecules, strictly depends on a non-enzymatic function of NIK

Previous studies have shown that p100 undergoes proteolytic processing to p52 only shortly after its synthesis, presumably before its incorporation into high-molecular-weight κBsome complexes^11^. To determine whether the non-enzymatic function of NIK promotes phosphorylation of newly synthesized p100, we first examined the relationship between the age of p100 molecules and the size of the complexes in which they reside.

Fractionation of tagged p100 complexes at different times after induction of p100 expression showed that newly synthesized p100 was predominantly present in complexes with apparent molecular masses below 300 kDa, whereas older p100 molecules, hereafter referred to as pre-existing p100, accumulated in larger assemblies with apparent molecular masses above 300 kDa (Fig. 2A).

**Figure 2.**
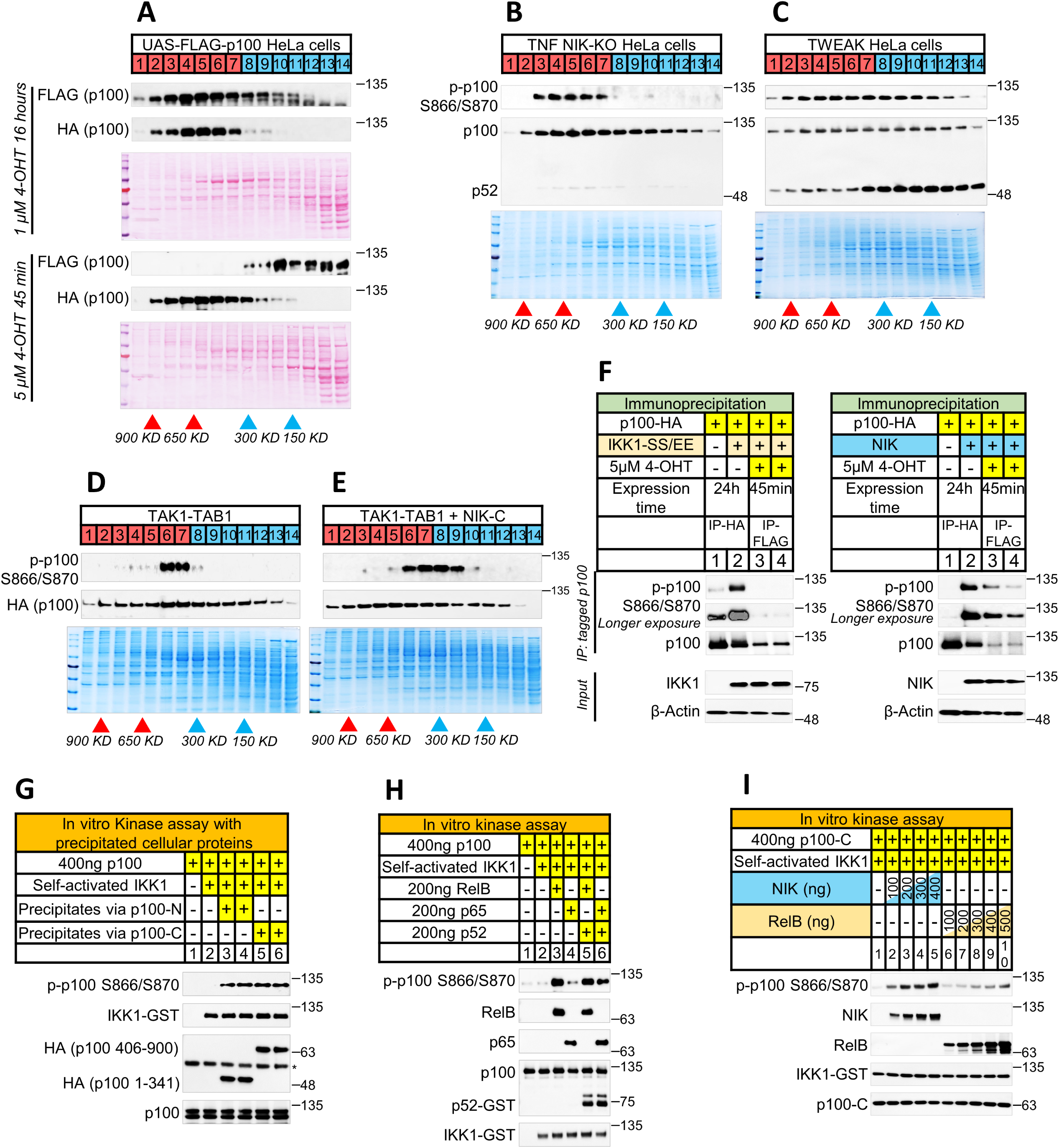
A non-enzymatic function of NIK is required for IKK1-mediated phosphorylation of newly synthesized p100, whereas NF-κB proteins facilitate phosphorylation of pre-existing p100 independently of NIK. A. Size-exclusion chromatography and immunoblot analysis of newly synthesized and pre-existing p100. FLAG-p100 expression was induced for 45 min or 16 h using the UAS/4-OHT system (see Methods), whereas HA-p100 was expressed constitutively. Because FLAG-p100 expression was substantially lower after 45 min of 4-OHT treatment than after 16h, the corresponding immunoblot was acquired using longer exposure time (40 s versus 10 s). Immunoblotting results are shown alongside the overall protein pattern in the samples, visualized by Ponceau S staining of the membrane. B. Size-exclusion chromatography and immunoblot analysis of TNF-induced phosphorylation of p100 in NIK-knockout HeLa cells. MG132 (10 μM) was added 1 h before fractionation. In this and all other panels showing size-exclusion chromatography results, except panel A, the overall protein pattern of the samples, shown alongside the immunoblot results, is visualized by Coomassie Blue staining. C. As in B, except that wild-type HeLa cells were stimulated with TWEAK. D. As in B, except that cells were co-transfected with plasmids encoding TAK1-TAB1. E. As in B, except cells were co-transfected with plasmids encoding TAK1-TAB1 together with NIK-C (residues 650–947). F. Analysis of phosphorylation of newly synthesized and pre-existing p100 in response to ectopic expression of NIK or IKK1-SS/EE. FLAG-p100 expression was induced by UAS/4-OHT system for 45 min and HA-p100 was ectopically expressed for 24 h. FLAG- and HA-tagged p100 molecules were recovered separately by anti-FLAG and anti-HA immunoprecipitation, respectively. G. In vitro phosphorylation of recombinant p100 by recombinant IKK1 in the presence or absence of immunoprecipitated p100-N (residues 1–341) or p100-C (residues 406–900) together with their associated cellular proteins (see Methods). Phosphorylation of recombinant p100 at S866/S870 was analyzed by immunoblotting. The asterisk indicates the immunoglobulin heavy chain of the antibody used for immunoprecipitation. H. In vitro phosphorylation of recombinant p100 by recombinant IKK1 in the presence or absence of recombinant RelB, p65, and p52 (see Methods). I. In vitro phosphorylation of recombinant p100-C (residues 406–900) by self-activated IKK1 in the presence of recombinant NIK or RelB.

We next asked whether NIK promotes phosphorylation of newly synthesized p100 molecules residing in low-molecular-weight complexes. To this end, we compared the size distribution of phosphorylated p100 in cells in which IKK1 was activated either by NIK or by NIK-independent mechanisms.

As shown in Fig. 2B, following TNF stimulation, phosphorylated p100 was detected almost exclusively in complexes larger than 300 kDa. The same pattern was observed in cells expressing TAK1-TAB1 (Fig. 2D) or IKK1-SS/EE (Fig. S2A). In contrast, in cells treated with TWEAK (Fig. 2C), which induces NIK accumulation, or in cells expressing ectopic NIK (Fig. S2B), a substantial fraction of phosphorylated p100 was detected in complexes smaller than 300 kDa.

Like the overall enhancement of p100 phosphorylation by NIK, phosphorylation of p100 in low-molecular-weight complexes did not require NIK kinase activity. Thus, as shown in Fig. 2D and 2E, co-expression of the C-terminal portion of NIK (residues 650–947; NIK-C) with TAK1–TAB1 enabled phosphorylation of p100 in these complexes. Consistently, co-expression of NIK-C with TAK1–TAB1 promoted accumulation of p52 (Fig. S2C), indicating that the non-enzymatic function of NIK was sufficient to promote p100 processing in this context.

These observations suggested that phosphorylation of pre-existing p100 molecules can occur independently of NIK, whereas phosphorylation of newly synthesized p100 requires a non-enzymatic function of NIK. We tested this prediction directly by monitoring phosphorylation of p100 at defined times after initiation of its synthesis. As shown in Fig. 2F, phosphorylation of p100 within 45 min of the initiation of its synthesis occurred only in cells expressing NIK (right panel). In contrast, phosphorylation detected 24 h after the initiation of p100 synthesis occurred both in cells expressing NIK and in cells expressing IKK1-SS/EE in the absence of NIK. Taken together, these findings indicate that the phosphorylation of newly synthesized p100 molecules by activated IKK1 strictly depends on a non-enzymatic function of NIK, whereas pre-existing p100 molecules can be phosphorylated by activated IKK1 independently of NIK.

As mentioned above, for reasons that remain unknown, only newly synthesized p100 molecules are competent for processing to p52 ^11^. The strict dependence of IKK1-mediated phosphorylation of such newly synthesized molecules on a non-enzymatic function of NIK provides a plausible explanation for why NIK-dependent phosphorylation, but not NIK-independent phosphorylation (of pre-existing p100), leads to p100 processing.

### Binding of other NF-κB proteins to p100 modestly enhances its phosphorylation by IKK1

The finding that activated IKK1 can phosphorylate pre-existing p100 within high-molecular-weight κBsome complexes even in the absence of NIK suggests that other proteins in these complexes may enhance phosphorylation of p100 by IKK1, albeit less efficiently than NIK. To test this possibility, we first examined whether proteins that associate with p100 in cells could enhance its phosphorylation by activated IKK1. We expressed fragments of p100 corresponding to its N-terminal and C-terminal portions (residues 1-341 and residues 406-900, respectively), isolated both fragments together with their associated cellular proteins, and assessed their ability to enhance IKK1-mediated phosphorylation of recombinant p100 in vitro. As shown in Fig. 2G, cellular proteins associated with both p100 fragments substantially enhanced phosphorylation of recombinant p100 by activated IKK1.

We therefore examined the effects of purified NF-κB proteins, which can associate with both the N-terminal and C-terminal portions of p100, on p100 phosphorylation in vitro. RelB alone and the combination of RelA (p65) and p52 markedly enhanced phosphorylation of p100 by activated IKK1, whereas neither RelA nor p52 alone did so (Figs. 2H and S2D). However, the magnitude of this enhancement was substantially lower than that produced by NIK (Fig. 2I).

This modest enhancement of p100 phosphorylation by NF-κB proteins is likely to account, at least in part, for the weak NIK-independent phosphorylation of pre-existing p100 molecules incorporated into NF-κB-containing κBsome complexes.

### Binding of NIK and IKK1 to the C-terminal region in p100 is required for S866/S870 phosphorylation and for its enhancement by NIK

NIK has previously been shown to bind both the Rel homology domain (RHD) in the N-terminal region of p100 and the ankyrin repeat (ANK) domain in its C-terminal region ^18^.

Deletion analysis revealed that IKK1, like NIK, interacts with both the N- and C-terminal regions of p100, but binds to sites distinct from those recognized by NIK (Figs. 3A, B and S3A). Consistent with a recent report ^19^, we identified an IKK1-binding site at the extreme N terminus of p100. This binding depended entirely on residues 1–33 and was further strengthened by downstream sequences (Fig. S3B). Association of IKK1 with this site and binding of NIK to the adjacent RHD were mutually enhanced (Figs. S3C and S3D). IKK1 also bound independently to a site in the C-terminal portion of p100 downstream of the ANK domain (residues 754–900) (Fig. S3A).

**Figure 3.**
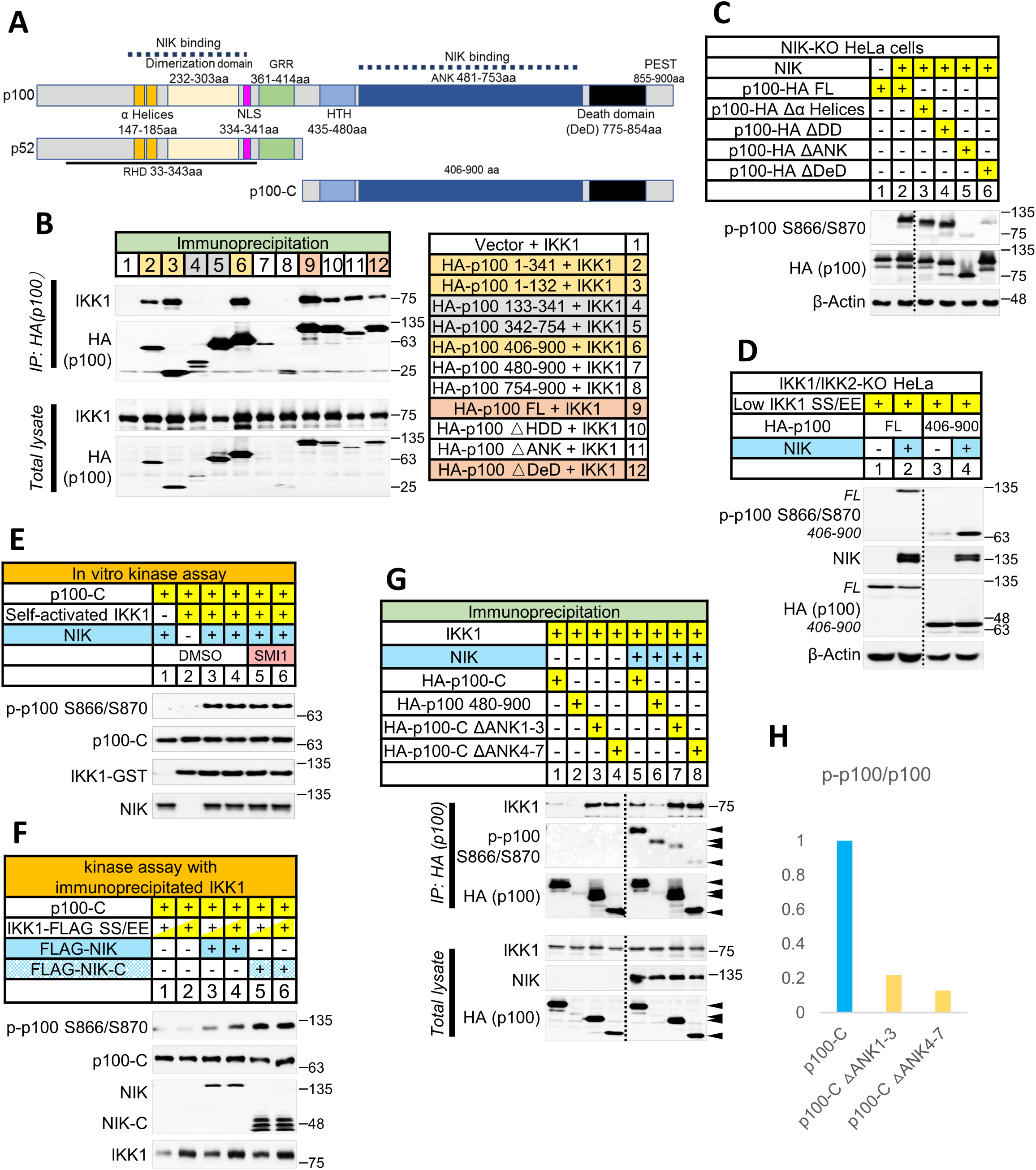
Phosphorylation of p100 by IKK1 and its enhancement by NIK require interactions with the C-terminal region of p100. A. Schematic representation of p100, p52, and p100-C. GRR, glycine-rich region; NLS, nuclear localization signal; RHD, Rel homology domain; HTH, helix-turn-helix region; ANK, ankyrin-repeat domain; DeD, death domain; PEST, proline-, glutamic acid-, serine-, and threonine-rich region. B. Co-immunoprecipitation of IKK1 with the indicated HA-tagged p100 fragments from lysates of NIK/p100-knockout HeLa cells co-transfected with plasmids encoding IKK1 and the indicated HA-tagged p100 fragments. C. Immunoblot analysis of phosphorylation of the indicated p100 deletion mutants in NIK-knockout HeLa cells co-transfected with plasmids encoding NIK and the indicated p100 deletion mutants. D. Immunoblot analysis of phosphorylation of full-length p100 and p100-C in IKK1/IKK2-knockout HeLa cells co-transfected with plasmids encoding NIK and IKK1-SS/EE (pBABE vector), together with a plasmid encoding either full-length p100 or p100-C. E. In vitro phosphorylation of recombinant p100-C by recombinant IKK1 in the presence or absence of recombinant NIK. SMI1 (200 nM) was added where indicated. F. In vitro phosphorylation of recombinant p100-C by FLAG-IKK1-SS/EE recovered by anti-FLAG immunoprecipitation from cell lysates containing FLAG-IKK1-SS/EE alone or from lysates mixed with lysates containing FLAG-NIK or FLAG-NIK-C before immunoprecipitation. FLAG-IKK1-SS/EE, FLAG-NIK, and FLAG-NIK-C were each expressed separately in IKK1-knockout HEK293T cells. G. Immunoblot analysis of IKK1-mediated phosphorylation of several p100 deletion mutants and their association with IKK1 in HeLa cells co-transfected with plasmids encoding IKK1 and NIK. MG132 (10 μM) was added 1 h before analysis. The following p100 mutants were used: p100 406-900 (p100-C); p100 480-900 (p100-C lacking the helix-turn-helix motif); p100 406-900 lacking residues 485–584 (designated as ΔANK1–3); and p100 406-900 lacking residues 596–752 (designated as ΔANK4–7). H. Bar chart showing the ratio of phosphorylated p100-C to total p100-C in (G).

We next asked which of these interactions is required for phosphorylation of S866/S870 in p100. Phosphorylation of p100 S866/S870 by IKK1, as well as its enhancement by NIK, occurred normally in cells expressing either a p100 mutant lacking the N-terminal IKK1-binding motif (Δ1–33; Fig. S3E) or mutants lacking the α-helical region or the dimerization domain within the NIK-binding RHD (Fig. 3C). In contrast, neither p100 S866/S870 phosphorylation nor its enhancement by NIK was observed in cells expressing p100 lacking either the ANK domain or the adjacent death domain (DeD) (Fig. 3C).

These findings indicate that phosphorylation of S866/S870 in p100 by activated IKK1 and its enhancement by NIK depend on interactions of both IKK1 and NIK with the C-terminal region of p100, whereas the N-terminal binding sites contribute little, if at all, to this process. These N-terminal interactions may serve other functions.

Consistent with this conclusion, NIK efficiently enhanced phosphorylation of p100 S866/S870 by IKK1-SS/EE in cells expressing only the C-terminal portion of p100 (residues 406–900; p100-C) (Fig. 3D). Similar enhancement was observed in a fully reconstituted in vitro assay using recombinant p100-C, recombinant NIK, and activated recombinant IKK1 (Fig. 3E). As expected for a non-enzymatic activity of NIK, this enhancement was unaffected by the NIK kinase inhibitor SMI1 (Fig. 3E). Moreover, such enhancement was also exerted by NIK-C (that was immunoprecipitated from cells together with IKK1-SS/EE, Fig. 3F).

NIK also efficiently promoted IKK1-mediated phosphorylation of p100 S866/S870 in cells expressing a C-terminal mutant of p100 lacking the helix-turn-helix region that mediates p100 dimerization ^14^ (p100 480-900, Fig. 3G). However, because the cellular levels of this mutant were extremely low, probably because the deletion of this motif compromises protein stability, we pursued the regulation of this phosphorylation using the full p100-C construct.

As with full-length p100, enhancement of p100-C phosphorylation by NIK depended on the ANK domain, to which NIK binds ^18^. Even partial deletions of this domain markedly reduced the ability of NIK to enhance phosphorylation of p100-C by either wild-type IKK1 (Fig. 3G and 3H) or IKK1-SS/EE (Fig. S3F). This reduction was not due to impaired binding of IKK1 to the mutant proteins. On the contrary, for reasons that remain unclear, these deletions markedly increased the binding of IKK1 to p100-C (Fig. 3G).

Together, these findings demonstrate that a non-enzymatic activity of NIK promotes IKK1-mediated phosphorylation of p100 through interactions entirely within the C-terminal region of p100.

### NIK restrains rather than enhances the kinase activity of phosphorylated IKK1

To determine whether NIK promotes p100 phosphorylation by further enhancing the kinase activity of phosphorylated IKK1, we first asked whether NIK remains associated with IKK1 after phosphorylating IKK1 and thereby modulates its activity.

Consistent with this possibility, immunoprecipitation of NIK from cell-lysates preferentially depleted phosphorylated IKK1 (Fig. S4A and B). Likewise, the phosphomimetic mutant IKK1-SS/EE displayed enhanced binding to NIK (Fig. S4C).

If this association augments the kinase activity of IKK1, it should also increase phosphorylation of substrates other than p100. To test this possibility, we examined phosphorylation of two established IKK substrates, IκBα and RelA (p65), in cells lacking IKK2 and expressing the constitutively active IKK1 mutants IKK1-SS/EE or IKK1-K170E (which is homologous to a constitutively active IKK2 mutant ^20^ and which we have found to phosphorylate various proteins more effectively than IKK1-SS/EE). IKK2-deficient cells were used because this kinase can also phosphorylate IκBα and p65 upon NIK activation independently of the kinase function of IKK1 ^18, 21^.

Contrary to this prediction, NIK did not enhance phosphorylation of either IκBα or p65. Instead, although it had no inhibitory effect in cells expressing IKK1-SS/EE together with IKK2 (Fig. 4A), NIK markedly inhibited phosphorylation of both substrates in cells expressing IKK1-SS/EE (Figs. 4A and B) or IKK1-K170E (Figs. S4D and E) in the absence of IKK2. This inhibition occurred under conditions in which NIK simultaneously enhanced phosphorylation of p100 by the same IKK1 mutants.

**Figure 4.**
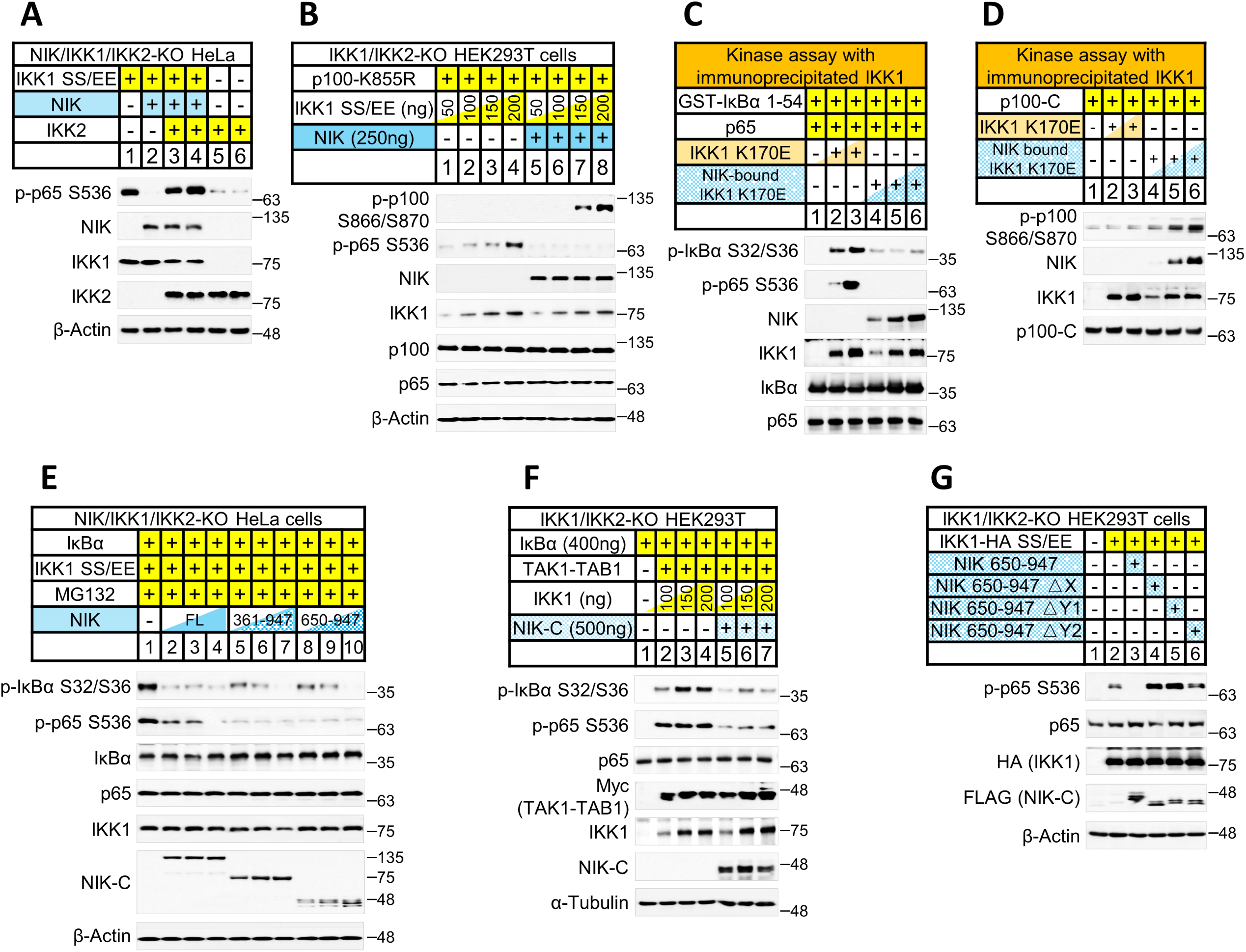
NIK remains associated with activated IKK1 and restrains its catalytic activity. A. Immunoblot analysis of p65 phosphorylation in NIK/IKK1/IKK2-knockout HeLa cells co-transfected with plasmids encoding IKK1-SS/EE and NIK, in the presence or absence of a plasmid encoding IKK2. B. Immunoblot analysis of p65 and p100 phosphorylation in IKK1/IKK2-knockout HEK293T cells co-transfected with a plasmid encoding p100 and increasing amounts of the plasmid encoding IKK1-SS/EE, in the presence or absence of a plasmid encoding NIK. C. In vitro phosphorylation of recombinant IκBα and p65 by immunoprecipitated IKK1-K170E, either alone or bound to NIK. IKK1-K170E was recovered by anti-FLAG immunoprecipitation from 30- or 60-μl lysates of cells transfected with a plasmid encoding FLAG-IKK1-K170E. NIK-bound IKK1-K170E was recovered by anti-HA immunoprecipitation from 60-, 90-, or 120-μl lysates of cells co-transfected with plasmids encoding FLAG-IKK1-K170E and HA-NIK. D. As in C, except that recombinant p100-C was used as the substrate. E. Immunoblot analysis of p65 and IκBα phosphorylation in NIK/IKK1/IKK2-knockout HeLa cells co-transfected with plasmids encoding IκBα and IKK1-SS/EE, together with increasing amounts (250, 500, or 750 ng) of plasmids encoding full-length NIK, NIK-361–947, or NIK-650–947. MG132 (10 μM) was added 1 h before analysis. F. Immunoblot analysis of p65 and IκBα phosphorylation in IKK1/IKK2-knockout HEK293T cells co-transfected with plasmids encoding TAK1-TAB1 and IκBα, together with increasing amounts of a plasmid encoding IKK1, in the presence or absence of a plasmid encoding NIK-C. MG132 (10 μM) was added 1 h before analysis. G. Immunoblot analysis of p65 phosphorylation in IKK1/IKK2-knockout HEK293T cells co-transfected with a plasmid encoding IKK1-SS/EE together with plasmids encoding NIK-C or the indicated NIK-C deletion mutants. ΔX, ΔY1, and ΔY2 denote deletion of residues 757–805, 847–892, and 893–947, respectively.

Both the inhibitory effect of NIK on IKK1-mediated phosphorylation of IκBα and p65 and its stimulatory effect on p100 phosphorylation were reproduced in cell-free assays (Figs. 4C and D). Notably, the inhibitory effect did not require the enzymatic activity of NIK. It was observed not only with full-length NIK and with an N-terminal deletion mutant (residues 361–947), but also with the non-enzymatic C-terminal portion of NIK (residues 650–947; NIK-C) (Fig. 4E). Likewise, expression of NIK-C effectively suppressed TAK1-TAB1-induced phosphorylation of both IκBα and p65 by IKK1 in IKK2-deficient cells (Fig. 4F).

The inhibitory activity required the X (residues 757–805) and Y (Y1: 847–892; Y2: 893–947) regions within NIK-C, both of which contribute to its binding to IKK1^18^. Deletion of either region abolished the inhibitory effect (Fig. 4G). The inhibition was also markedly reduced by an X region mutant carrying three leucine-to-alanine substitutions, which weakens NIK binding to the kinase domain of IKK1 without affecting NIK-mediated phosphorylation of IKK1^18^ (Fig. S4F). Thus, the interaction required for inhibition of activated IKK1 is structurally more stringent than that required for activation of IKK1 by phosphorylation.

Taken together, these findings show that the enhancement of IKK1-mediated p100 phosphorylation by NIK does not result from augmentation of the kinase activity of IKK1. On the contrary, association of NIK with activated IKK1 suppresses the ability of the latter to phosphorylate substrate proteins. This finding strongly suggests that the enhancement of IKK1-mediated p100 phosphorylation by NIK reflects an effect of NIK on the substrate rather than on the kinase.

### Single-molecule FRET spectroscopy reveals that NIK induces a conformational change in p100-C

Using single-molecule Förster resonance energy transfer (smFRET) spectroscopy, we sought to determine how p100-C is affected by NIK binding. We took advantage of the five cysteine residues in p100-C: C891, located within the distal segment of the PEST region (residues 871–900), downstream of the IKK1 phosphorylation sites S866/S870, and four upstream cysteines (C432, C615, C703, and C738). To assess the proximity of the distal PEST region to these upstream sites, we generated four recombinant p100-C FRET reporters in which three of the four upstream cysteines were replaced with serines, leaving one upstream cysteine and C891 available for fluorescent labeling. These reporters were designated C432–C891, C615–C891, C703–C891, and C738–C891.

Three of these reporters bound NIK as effectively as wild-type p100-C (Fig. S5A). Because NIK binding to the C738–C891 reporter was substantially reduced, this reporter was not analyzed further. The remaining three reporter proteins were labeled with Alexa Fluor 488 and Alexa Fluor 594 and analyzed as individual molecules freely diffusing through a focused laser beam (Fig. 5A). Fluorescence bursts were identified and analyzed to generate FRET efficiency distributions.

**Figure 5.**
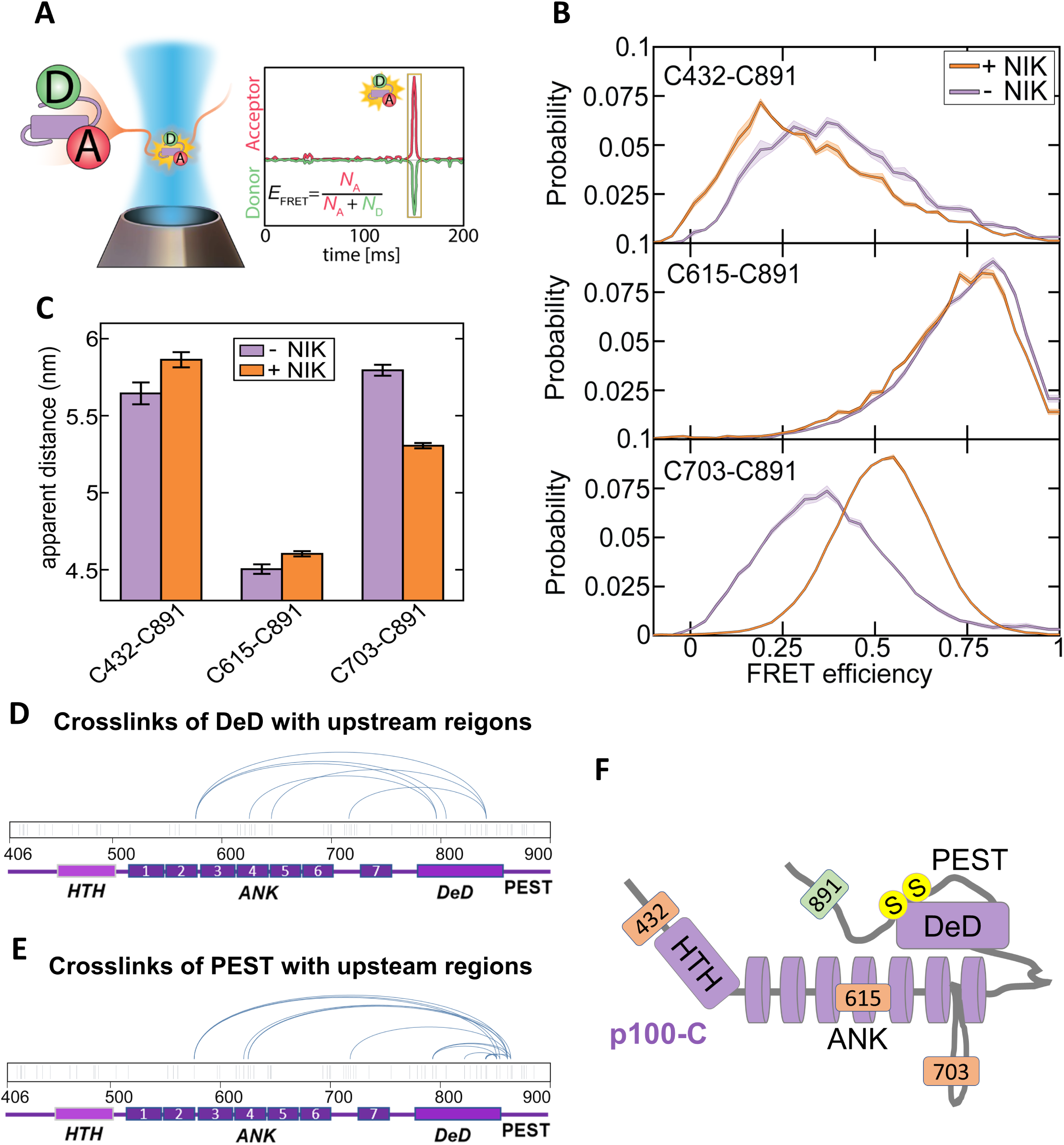
The C-terminal region of p100 adopts a folded-back conformation which is altered upon binding of NIK to it. **A.** Schematic representation of smFRET experiments. p100-C molecules labeled with donor (D) and acceptor (A) dyes generated bursts of fluorescence photons as individual molecules diffused through a focused laser beam. D (green) and A (red) photons originating from each fluorescence burst were extracted from the time traces to determine the number of acceptor (N_A_) and donor (N_D_) photons. The ratio of acceptor photons to the total number of detected photons for each fluorescence burst defines the FRET efficiency (E_FRET_), which reports on the relative proximity of the two dyes. **B.** FRET efficiency distributions for the indicated p100-C reporters in the absence (purple) or presence (orange) of NIK. Addition of NIK produced a marked increase in FRET efficiency for the C703–C891 reporter, a modest decrease for C432–C891, and little change for C615–C891. Solid lines represent the mean of 4 independent measurements, and shaded regions indicate the standard error of the mean. **C.** Apparent inter-dye distances derived from smFRET measurements for the C432–C891, C615–C891, and C703–C891 reporters in the absence or presence of NIK. Error bars indicate the standard error of the mean from 4 independent measurements. **D** and **E**. Arc maps of DHSO cross-links connecting the death domain (DeD; residues 754–847) (D) or proximal PEST region (residues 850–870) (E) to upstream regions of p100-C. Each arc line connects the identified cross-linking sites. Short vertical ticks indicate all Asp and Glu residues within p100-C, representing potential DHSO cross-linking sites. Cross-links involving HTH-adjacent regions are omitted from these maps but retained in the complete cross-link tables. The domain organization of p100-C is shown below each map. **F.** Schematic model of the conformational organization of p100-C supported by the smFRET and XL-MS data. The model illustrates the proximity of the DeD and proximal PEST region to the ANK domain without assigning a unique conformation to the flexible PEST region.

As shown in Fig. 5B and C, addition of NIK markedly shifted the FRET efficiency distribution of the C703–C891 reporter, increasing the peak FRET efficiency from approximately 0.35 to 0.53. This change indicates a reduction in the apparent distance between C703 and C891. In contrast, addition of NIK to the C432–C891 reporter resulted in a modest decrease in peak FRET efficiency, whereas little change was observed with the C615–C891 reporter.

Control experiments showed that an unrelated protein did not reproduce the observed changes in FRET efficiency (Fig. S5B), excluding nonspecific protein interactions or molecular crowding as explanations for these changes. Changes in dye rotational freedom were also excluded as a possible explanation (Fig. S5C).

These findings indicate that NIK binding to p100 dictates a conformational change in the latter. The differential responses of the three FRET reporters suggest that NIK induces a selective rearrangement of p100-C rather than uniform compaction or expansion.

### Both single-molecule FRET spectroscopy and cross-linking mass spectrometry suggest a folded-back conformation of p100-C

Because all three FRET reporters shared C891 as a common reference site, their FRET efficiencies allowed us to compare the relative proximity of the three upstream labeling sites to C891. We therefore used the average FRET efficiency values to calculate apparent inter-dye distances from the corrected FRET efficiencies (Fig. 5C). The C615–C891 pair displayed the shortest apparent distance, followed by C703–C891. This suggests that the distal PEST region is not simply localized near the C-terminal end of the ANK domain, but instead preferentially folds back toward its central portion. The apparent distance between C432 and C891 was comparable to that between C703 and C891, indicating that the N-terminal region of p100-C also lies relatively close to the distal PEST segment. The FRET efficiency distributions of all three reporters were two- to fourfold broader than expected from photon shot noise alone, suggesting that this spatial organization accommodates a range of conformations.

Together, these observations support a folded-back organization in which the distal segment of the PEST region samples a spatially restricted but heterogeneous conformational ensemble relative to upstream regions of p100-C.

Having obtained evidence for a folded-back organization of the PEST region and a conformational rearrangement of p100-C upon NIK binding, we next sought to determine how the DeD and PEST regions are positioned relative to upstream regions of p100-C. To address this question, we used cross-linking mass spectrometry (XL-MS) to identify spatial proximities involving these regions. We employed two distinct cross-linking strategies: DHSO in combination with DMTMM, which links two carboxyl-containing residues through a defined spacer ^22^, and DMTMM alone, which mediates coupling of activated carboxyl groups to amino groups of lysine and, under certain conditions, to the hydroxyl groups of serine, threonine, and tyrosine residues ^23,24^.

DHSO cross-linking identified reproducible links between the DeD and upstream regions of the ANK domain (Fig. 5D), supporting an organization in which the DeD folds back toward the ankyrin repeats. Cross-links involving the PEST region were confined to its proximal segment (residues 850–870) and connected this segment to upstream regions within the ANK domain (Fig. 5E); no cross-links involving its distal segment were detected. DMTMM cross-linking provided additional evidence for these spatial proximities (Fig. S5F and G; Table S1). Together, these cross-linking results complement the smFRET measurements and support a folded-back organization of p100-C (Fig. 5F). In this organization, the close proximity of the DeD and the proximal PEST segment to upstream ANK regions may constrain the C891-containing distal segment of the PEST region to a spatially restricted but heterogeneous conformational ensemble.

As shown above, NIK differentially affected the three FRET reporters. The pronounced change in FRET efficiency observed with the C703–C891 reporter, compared with the modest change with the C432–C891 reporter and little change with the C615–C891 reporter (Fig. 5 B and C), suggests that NIK preferentially repositions the C703-containing region relative to the C891-containing distal PEST segment. C703 lies in the linker between ANK6 and ANK7, within the ANK5–7 region to which the DeD and proximal PEST segment predominantly cross-linked.

This correspondence suggests that the NIK-induced rearrangement may involve changes in the relative positioning of the DeD and proximal PEST segment with respect to the ANK domain. Although these experiments do not define the precise nature of this rearrangement or directly measure the accessibility of the phosphorylation sites, they support a model in which the folded-back organization of p100-C limits access to S866/S870, whereas NIK-induced rearrangement may increase their accessibility to IKK1, providing a possible structural basis for the NIK-dependent enhancement of their phosphorylation.

## DISCUSSION

The primary goal of this study was to clarify the mechanism by which NIK—the kinase that initiates the non-canonical NF-κB pathway by phosphorylating, and thus activating, IKK1—also imposes specificity on the function of the latter.

NIK functions downstream of a restricted set of biological stimuli, primarily signals from certain TNF family receptors involved in the regulation of adaptive immunity. In contrast, IKK1 can be activated by multiple upstream kinases and phosphorylate a variety of substrate proteins, thereby eliciting diverse and functionally distinct responses. The functional specificity of the non-canonical NF-κB pathway therefore requires a mechanism that restricts that phosphorylation of p100 that leads to its processing to stimuli that activate IKK1 through NIK, rather than allowing these events to occur whenever IKK1 is activated.

For some time, it was thought that IKK1 molecules activated in the absence of NIK are completely incapable of phosphorylating S866/S870 in p100. However, both the present study and a recent independent study ^17^ showed that IKK1 activated through the canonical NF-κB pathway—which is initiated primarily by ligands that regulate innate immunity and in which IKK1 is activated by kinases other than NIK, including TAK1—can also phosphorylate these residues. This phosphorylation, however, is less efficient than that mediated downstream of NIK and, unlike the latter, does not promote processing of p100 to p52.

Our findings provide an initial clue as to why only phosphorylation of p100 by IKK1 activated by NIK promotes its proteolytic processing. Consistent with the restriction of p100 processing to newly synthesized molecules ^11^, we show that IKK1-mediated phosphorylation of newly synthesized p100 occurs only in cells expressing NIK, whereas the modest phosphorylation of p100 by IKK1 in the absence of NIK is restricted to p100 molecules that have already assembled into high-molecular-weight κBsomes (Fig. 6A). Thus, phosphorylation of p100 that leads to p52 generation is restricted to signaling downstream of NIK and adaptive immune regulators, whereas phosphorylation induced downstream of innate immune signaling affects only p100 molecules that reside in κBsomes. This distinction is consistent with the differing biological roles of these signaling pathways.

**Figure 6.**
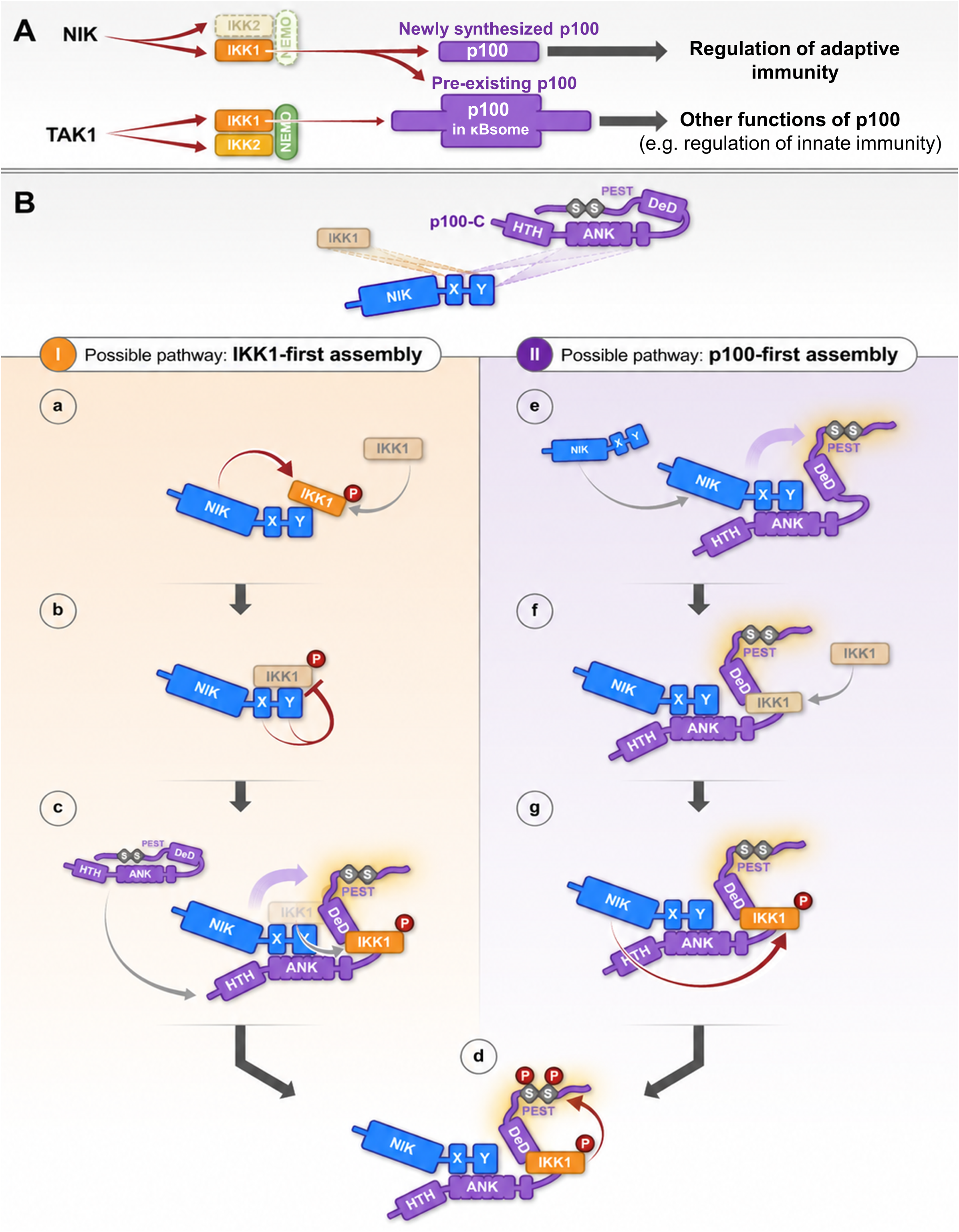
Diagrammatic illustration of the mechanisms proposed to underlie the non-enzymatic ‘licensing’ by NIK of IKK1-mediated phosphorylation of p100. A. The unique features of p100 phosphorylation downstream of NIK Comparison of the phosphorylation of serine residues 866 and 870 in p100 by IKK1 following its activation by either NIK or another MAP3K, TAK1. Although in both cases activated IKK1 can phosphorylate p100, phosphorylation downstream of NIK is much more efficient. Moreover, whereas IKK1-mediated phosphorylation of pre-existing p100 assembled into κBsomes can be observed in both cases, phosphorylation of newly synthesized p100 molecules is observed exclusively in cells expressing NIK. Whether NIK-mediated enhancement is restricted to newly synthesized p100 or also extends to the pre-existing p100 pool remains uncertain. **B.** Two possible sequences of events by which NIK facilitates IKK1-mediated phosphorylation of p100 through a non-enzymatic mechanism The illustration of NIK, IKK1, and the C-terminal portion of p100 (p100-C) at the top of the figure highlights the fact that both IKK1 and the ankyrin-repeat domain of p100 (ANK) can bind NIK by associating with the same two structured domains in its C-terminal region, the X and Y domains. p100-C adopts a compact, folded-back conformation when not associated with NIK.

Our finding that several NF-κB proteins (e.g., RelB or a combination of RelA and p52) modestly enhance IKK1-mediated phosphorylation of p100 in the absence of NIK provides a possible explanation for how p100 molecules incorporated into κBsomes become susceptible to limited NIK-independent phosphorylation by activated IKK1. Why such NIK-independent phosphorylation of p100 by IKK1 does not similarly affect newly synthesized p100, why only phosphorylation of newly synthesized p100 leads to its processing to p52, and what functional role is served by IKK1-mediated phosphorylation of p100 incorporated into κBsomes remain to be clarified. It also remains to be established whether differences in the extent of NIK dependence of p100 phosphorylation indeed relate to the age of p100 molecules or to their assembly into low-versus high-molecular-weight complexes.

By what mechanism does NIK, after phosphorylating the serine residues in the activation loop of IKK1, further enhance IKK1-mediated phosphorylation of p100?

We found that the NIK-mediated enhancement of p100 phosphorylation can be reconstituted in vitro using recombinant NIK, IKK1, and p100-C (residues 406–900) and that the C-terminal region of NIK (residues 650–947), which lacks the kinase domain, was sufficient for this effect. Together, these findings teach that this enhancement is exerted by NIK non-enzymatically, via some direct effect that it has either on IKK1 or on p100-C.

We considered the possibility that NIK, after phosphorylating IKK1, remains bound to it and further enhances its activity. To our surprise, we found that although phosphorylated IKK1 indeed binds avidly to NIK, this association does not further enhance IKK1 activity but instead suppresses it. Notably, the C-terminal region of NIK is likewise sufficient for this inhibitory effect. This suppression is selective: NIK inhibits IKK1-mediated phosphorylation of two other substrate proteins, IκBα and RelA (p65), whereas phosphorylation of p100 is spared from this inhibition and is ultimately enhanced in the presence of NIK. Fig. 6B presents two possible mechanisms by which this selectivity might be achieved. Because the same structural motifs in the C terminus of NIK mediate binding to both IKK1 and p100 ^18^, binding of p100 to NIK could disrupt or weaken the inhibitory NIK–IKK1 interaction through competition for the X and Y domains of NIK, thereby relieving the inhibitory effect that NIK exerts on IKK1 activity (Fig. 6B, I). Alternatively, because the IKK1-binding site in p100-C is distinct from the NIK-binding site, it is possible that many—or even all—of the IKK1 molecules that phosphorylate p100 are recruited to p100 before being phosphorylated by NIK and therefore never become subject to NIK-mediated inhibition (Fig. 6B, II).

Regardless of which of these mechanisms predominates, mere relief of NIK-mediated inhibition of IKK1 upon NIK binding to p100 cannot account for the enhanced p100 phosphorylation. Having established that the non-enzymatic function of NIK does not involve further enhancement of IKK1 kinase activity, we sought to gain insight into how NIK affects the conformation of p100 upon binding. Because both p100-C and the region of NIK that binds to it contain multiple flexible regions, several of which are structurally unresolved, we employed single-molecule FRET (smFRET) spectroscopy and XL-MS. These analyses revealed that p100-C adopts a compact, folded-back conformation in which the DeD- and PEST-containing region folds back toward and lies in close proximity to upstream regions. This structural arrangement provides a plausible explanation for the limited accessibility of the S866/S870-containing region to IKK1. smFRET analysis further revealed that NIK binding alters the relative position of the PEST region and upstream ANK-containing region, suggesting that NIK perturbs this folded-back conformation and thereby potentially increases the accessibility of the S866/S870-containing region to IKK1 (Fig. 6B).

Our findings reveal that NIK does not merely act as an activating kinase upstream of IKK1. By transiently restraining the function of phosphorylated IKK1 while directly modulating p100, NIK acts as a substrate-specifying licensing factor. We previously showed that IKK1 can perform a non-catalytic scaffolding function to facilitate NIK-mediated activation of IKK2 and canonical NF-κB proteins ^18^. Here, we show that, in the non-canonical pathway, NIK similarly couples its enzymatic activity to a distinct non-enzymatic function.

These findings provide a conceptually distinct example of signaling regulation. Many signaling pathways regulate substrate specificity independently of activation of the initiating enzyme, and many signaling proteins possess both catalytic and non-catalytic functions. However, to our knowledge, our findings provide the first example of a signaling mechanism in which a protein kinase, in addition to initiating a signaling cascade enzymatically, directly acts on a downstream substrate of the cascade as a non-enzymatic target-licensing factor. The strict dependence of phosphorylation of newly synthesized p100 on this licensing function of NIK imposes specificity on IKK1, ensuring that p100 processing and the resulting gene activation by p52-containing NF-κB dimers occur only in response to NIK-inducing stimuli.

These findings also have important implications for the attempts to develop novel means of therapy. Use of NIK-inhibitory drugs for therapy of various autoimmune diseases and cancer is now being explored ^25,26^. However, our findings suggest that pharmacological inhibition of NIK kinase activity may not fully recapitulate the effects of complete NIK deficiency, as such NIK-inhibitory drugs would not be expected to disrupt the non-enzymatic interaction between NIK and p100. This distinction may be particularly relevant under pathological conditions in which TNF and other canonical pathway stimuli activate IKK1 independently of NIK, while non-canonical signaling sustains the NIK–p100 licensing interaction. Targeting the interaction underlying this licensing function may therefore offer an alternative strategy for limiting pathological p100 processing.

## LIMITATION OF THE STUDY

Although our findings clearly show that NIK both enhances IKK1-mediated phosphorylation of p100 and induces a conformational change in p100, they do not establish a causal relationship between these two effects.

Moreover, how NIK alters p100 conformation and how this conformational change might promote the phosphorylation of p100 remain to be determined.

The present study also does not resolve the order of assembly of the NIK–IKK1–p100 complex. One possibility is that p100 binding disrupts or weakens the inhibitory NIK–IKK1 interaction through competition for NIK binding, whereas another is that the complex assembles sequentially, with NIK first binding p100 and subsequently recruiting IKK1 (as illustrated in Fig. 6).

NIK and IKK1 both bind to sites in the C-terminal, IκB-like region of p100 and in its N-terminal region. Our findings indicate that only their interactions with the C-terminal region contribute to IKK1-mediated phosphorylation of S866/S870 and its enhancement by NIK. The functions served by NIK and IKK1 binding to the N-terminal region remain to be determined.

We found that NIK-independent phosphorylation of p100 by IKK1, which does not lead to p100 processing to p52, is restricted to p100 molecules incorporated into κBsomes. The functional significance of this phosphorylation remains unclear.

In addition, although we found that several NF-κB proteins partially enhance p100 phosphorylation, other p100-binding proteins may also contribute to the enhancement of IKK1-mediated phosphorylation of p100 within κBsomes.

Finally, the physiological significance of NIK-mediated inhibition of IKK1 activity toward substrates other than p100 remains to be determined.

### Pathway I (‘IKK1-first’ assembly)

**a.** When encountering IKK1, NIK binds to it and phosphorylates the two serine residues S176 and S180 in its activation loop.
**b.** S176/S180-phosphorylated IKK1 binds more avidly to the X and Y domains of NIK. This binding inhibits IKK1 function. Thus, paradoxically, although S176/S180 phosphorylation increases the intrinsic kinase activity of IKK1, phosphorylated IKK1 remains functionally restrained when avidly associated with NIK.
**c.** When p100 (or p100-C) encounters the NIK–IKK1 complex, it may compete with IKK1 for binding to the X and Y domains of NIK, thereby disrupting or weakening the inhibitory NIK–IKK1 interaction and relieving inhibition of IKK1 kinase activity. NIK and IKK1 then engage distinct sites in p100-C—NIK binding to ANK and IKK1 to a C-terminal site downstream of ANK. Binding of NIK to ANK in p100-C induces a conformational change in p100-C that likely increases the accessibility of the IKK1 substrate residues in the PEST region (S866/S870).
**d.** The active IKK1 molecule bound to p100-C phosphorylates S866/S870 in the PEST region, which has become more accessible following NIK binding to ANK, as proposed in **c.**

### Pathway II (‘p100-first’ assembly)

**e.** NIK binds to the ANK domain in p100-C. NIK binding induces a conformational change in p100-C that likely increases the accessibility of the PEST region containing S866/S870.
**f.** IKK1 binds to p100-C at a site C-terminal to the NIK-binding site.
**g.** The NIK molecule bound to p100-C phosphorylates the nearby IKK1 molecule. Because NIK is already engaged with p100, phosphorylated IKK1 avoids the aforementioned avid inhibitory interaction with the X and Y domains of NIK. The activated IKK1 molecule can then phosphorylate S866/S870 in p100-C, as shown in **d**.

## Supporting information

Suppemental figures

## ACKNOWLEDGMENT

We wish to thank Professor Rony Seger from the Weizmann Institute of Science (WIS) for his advice on the in vitro protein-kinase reactions. We thank Drs. Yoav Peleg and Shira Albeck from the Structural Proteomics Unit, Life Sciences Core Facilities, WIS for the production and purification of recombinant p100-C and p100-C mutants, Dr. Guy Zoltsman from the WIS for assistance with protein purification, Tatiana Shalevich for maintaining cultured cells, and both Professors Rony Seger and Michael Walker from the WIS for helpful advice on the manuscript. This research was fully funded by the own resources of the lead contact of this publication.

## AUTHOR CONTRIBUTIONS

X.L. and D.W. were responsible for conception of most of the project and for data analysis. X.L. performed most of the experiments. D.S. and G.H. were responsible for conception of the smFRET analysis, and D. S. did all the experiments associated with this analysis. D.M. and D.R. contributed to the XL-MS experiments presented in Fig. 5 and Fig. S5. K.B. generated DNA constructs. Mike G., C.A. and T. W. contributed to experiments associated with Figs 1, 3 and 4. Moshe G. contributed to experiments associated with Fig. 2, and D.W. supervised the study.

## COMPETING INTEREST DISCLOSURE STATEMENT

The authors declare no competing interests.

## SUPPLEMENTARY INFORMATION

Figure S1, Figure S2, Figure S3, Figure S4, Figure S5.

Table S1. Complete list of XL-MS crosslinks identified in p100-C.

Table S2. gRNA seed sequences used in this study.

## MATERIALS AND METHODS

### Cell culture and transfection

HeLa cervical adenocarcinoma cells and HEK293T cells were cultured in Dulbecco’s Modified Eagle Medium (DMEM) supplemented with 10% fetal bovine serum (FBS), 100 U/ml penicillin, and 100 μg/ml streptomycin. Cells were transfected 18–24 h after seeding at 50%–80% confluency using jetPEI (Polyplus) according to the manufacturer’s instructions.

### Plasmid construction

Human p100, IKK1, and IKK2 cDNAs were purchased from Invitrogen, whereas human NIK cDNA was cloned from a human B-cell cDNA library. Coding sequences were amplified by PCR with the indicated tags or adaptors and cloned into the appropriate expression vectors using standard restriction enzyme-mediated ligation. Deletion and point mutants were generated using a QuikChange Site-Directed Mutagenesis Kit (Stratagene) according to the manufacturer’s instructions. All plasmid constructs were verified by DNA sequencing.

### Generation and selection of CRISPR knock-out cell lines

Cells were seeded into 6-well plates at approximately 40% confluency and transfected with PX459 plasmids encoding the indicated guide RNAs. 18–24 h after transfection, cells were subjected to puromycin selection until all non-transfected control cells were eliminated. Puromycin was used at 1 μg/ml for HEK293T cells and 0.5 μg/ml for HeLa cells. Surviving cells were expanded and single-cell clones were isolated by limiting dilution in 96-well plates using growth medium supplemented with 1 mM sodium pyruvate (Biological Industries, Cat# 03-042-1B) and non-essential amino acids (Biological Industries, Cat# 01-340-1B). Knockout clones were validated by immunoblotting. Guide RNAs (Table S2) were designed using an online CRISPR design tool, synthesized by Integrated DNA Technologies, and cloned into PX459 (Addgene plasmid #48139).

### Lentivirus production and transduction

For lentivirus production, pF5UAS-NIK, pF5UAS-p100, or pFUGEV16 was separately transfected into HEK293T cells together with the packaging plasmids pLP1, pLP2, and pLP/VSVG (Invitrogen). Viral supernatants were collected 48 h after transfection and passed through a 0.45-μm sterile syringe filter (Cytiva). HeLa cells were first transduced with GEV16 lentivirus in the presence of 8 μg/ml Polybrene. 8 h after transduction, 1.5 volumes of fresh culture medium were added. Cells were passaged 48 h later and selected in growth medium containing 50 μg/ml hygromycin. The resulting GEV16-expressing cells were subsequently transduced with UAS-NIK or UAS-p100 lentivirus under the same conditions and selected in growth medium containing 0.5 μg/ml puromycin.

### Inducible expression of NIK or p100

Stable HeLa cell lines expressing inducible NIK or p100 were generated in NIK-knockout or p100-knockout cells, respectively, using the GEV16/pF5UAS lentiviral expression system. NIK or p100 expression was induced by treatment with 4-hydroxytamoxifen (4-OHT).

### Cell lysis

Unless otherwise indicated, cells were washed twice with ice-cold PBS and then lysed for 20 min at 4°C in HEPES lysis buffer containing NP40 (30 mM HEPES, pH 7.4, 150 mM NaCl, 1 mM EGTA, 100 mM NaF, 1 mM Na VO, 10 mM β-glycerophosphate, 5 % glycerol, 0.5 % NP40, 0.5 % sodium deoxycholate, and cOmplete™ Protease Inhibitor Cocktail). Protein concentration in the cell lysates was measured using the Pierce 660 nm Protein Assay (Thermo Fisher Scientific).

### Antibodies, chemicals and other reagents

Unless otherwise indicated, primary antibodies were used for immunoblotting at a dilution of 1 to 1,000. In this study we employed antibodies against NIK (4994, cell signaling), p52/p100 (05-361, Merck Millipore), IKK1 (2682, cell signaling; 556531, BD bioscience), IKK2 (2684, cell signaling; 551920, BD bioscience), p-IKKs (2697, cell signaling), p-p100 S866/S870 (4810, cell signaling), p-IκBα (9246, cell signaling, sc-8404, SANTA-CRUZ), IκBα (ab32518, Abcam), p-p65 S536 (3033, cell signaling), p65 (8242, cell signaling), p50/p105 (3035; 13586, cell signaling), RelB (4922, cell signaling), α-Tubulin (ab7291, Abcam), β-Actin (A5441, Sigma), HA (ab9110, Abcam), FLAG (F3165, Sigma).

We also employed Anti-FLAG agarose (A-2220, Sigma), Anti-HA agarose (A-2095 Sigma) and Protein G agarose (GE 17-0618-01 Cytiva), SMI1 (hy-112433, MedChemExpress), DHSO (909599, Sigma), DMTMM (74104-1G-F, Sigma), MG132 (M7449, Sigma), Puromycin (ant-pr-1, Invitrogen), G418 disulfate (A1720, Sigma), Recombinant TWEAK (310-06-100, Peprotech), Recombinant TNFα (300-01A-10, Peprotech), Recombinant NIK (1065H, Creative BioMart), Recombinant IKK1 (AB102103, Abcam), recombinant p100 (TP313313, OriGene), recombinant p100-C (home-made), recombinant RelB (R30-30H, SinoBiological), recombinant p52 (N13-31G, SinoBiological), recombinant p65 (TP320780, OriGene), and recombinant IκBα (PKSH030926, Elabscience).

### Size exclusion chromatography

Cell lysates were prepared from one 150 mm cell culture dish (Nunc™ 168381) using HEPES lysis buffer as described above, clarified by centrifugation, and normalized to a total protein concentration of 4–8 mg/ml.

Size exclusion chromatography was carried out on an ÄKTA pure system (GE Healthcare). A 500 μl aliquot of each lysate was loaded onto a Superose 6 Increase 10/300 GL column (GE29-0915-96, Sigma) and fractionated in PBS containing 150 mM NaCl at a flow rate of 0.5 ml/min. Fractions (350 μl each) were collected, and 10–25 μl of the indicated fractions were analyzed by immunoblotting or Coomassie Brilliant Blue staining.

The apparent molecular weight range of proteins eluting in each fraction was estimated from a calibration curve generated using commercially available molecular weight standards, including carbonic anhydrase, bovine serum albumin, β-amylase, bovine IgG, and thyroglobulin.

### In vitro kinase assay

Recombinant IKK1 was used for in vitro phosphorylation of p100. To activate recombinant IKK1, the protein (10 ng/μl) was incubated in kinase buffer containing 1 mM ATP at 30 °C for 1 h. Activated recombinant IKK1 (100 ng) was then added to kinase buffer containing 1 mM ATP together with recombinant p100 (400 ng; full-length or residues 406–900) in the presence or absence of immunoprecipitated p100 fragments together with their associated cellular proteins, or recombinant NIK, RelB, RelA (p65) or p52 (200–400 ng each, as indicated). Reactions were performed in a final volume of 40 μl at 30 °C for 45 min and stopped by adding 40 μl of 2 × LDS sample buffer (Invitrogen, Cat# NP0007). Where indicated, the NIK inhibitor SMI1 (200 nM) was included to inhibit NIK kinase activity.

### IP kinase assay

IKK1-SS/EE or IKK1-K170E was immunoprecipitated either alone or together with NIK or NIK-C. The immunoprecipitates were washed twice with ice-cold PBS and once with kinase buffer (9802, Cell signaling). Recombinant p100, p100-C, IκBα, or p65 (400 ng for each, as indicated) was incubated with the immunoprecipitated IKK1-SS/EE or IKK1-K170E in kinase buffer containing 1 mM ATP at 30 °C for 45 min. Reactions were performed in a final volume of 40 μl and stopped by adding 40 μl of 2× LDS sample buffer.

### Recombinant p100-C preparation

His-tagged recombinant p100-C (wild-type and various mutants) was expressed in *E. coli* and first purified by Ni²-affinity chromatography. Bound protein was eluted with imidazole, and fractions containing p100-C were identified by SDS-PAGE and pooled. The pooled material was subsequently subjected to size-exclusion chromatography on a Superdex 200 column. Fractions containing the major dimeric species were pooled, whereas earlier-eluting high-molecular-weight aggregates were excluded. The final preparation was analyzed by SDS-PAGE and used for subsequent experiments.

Consistent with the reported strong dimerization propensity of p100-C, the SEC profile showed a predominant dimeric species (90% of the pooled protein) and only a minor portion of monomer. The oligomeric state of purified recombinant p100-C was characterized by mass photometry at concentrations of 1, 10, 20, and 30 nM. The proportion of dimeric p100-C decreased markedly with decreasing protein concentration, with monomeric species accounting for more than 80% of the detected particles at 1 nM (Fig. S5H).

### Mass photometry

Mass photometry measurements were performed using a OneMP mass photometer (Refeyn Ltd., Oxford, UK). Microscope coverslips (No. 1.5, 24 × 50 mm; cat. no. 0107222, Marienfeld) were cleaned by sequential sonication in 50% HPLC-grade isopropanol in Milli-Q water and Milli-Q water (5 min each), followed by drying under a stream of nitrogen. Gaskets (MP-CON-21022, Refeyn) were placed at the center of each coverslip, with each well used for a single measurement. Filtered buffer was first added to the well to establish and stabilize the focal position using the droplet-dilution mode. Purified recombinant p100-C was measured at final concentrations of 1, 10, 20, and 30 nM. For each acquisition, the protein sample was added to the buffer and mixed several times, and data were collected for 2 min at room temperature using the regular window size. Monomeric and dimeric BSA (Sigma), bovine IgG (Sigma), and MassFerence P1 calibrant (Refeyn) were used for mass calibration. Data were acquired using AcquireMP (Refeyn, 2025 R1.2) and analyzed using DiscoverMP (Refeyn, 2025 vR1).

### Immunoprecipitation and immunoblotting

For the immunoprecipitation experiments, cells were lysed in RIPA buffer (50 mM Tris-HCl, pH 7.4, 150 mM NaCl, 1% Triton X-100, 0.5% sodium deoxycholate, 0.1% SDS, and cOmplete™ Protease Inhibitor Cocktail). Cell lysates were pre-cleared with Protein G-agarose for at least 1 h at 4 °C and clarified by centrifugation at 13, 000 × g for 10 min. The lysates were then incubated with the indicated primary antibodies for at least 4 h at 4 °C, followed by incubation with Protein G-agarose for 1 h. Where indicated, anti-HA or anti-FLAG agarose beads were used directly for immunoprecipitation. The beads were washed three times with the lysis buffer and resuspended in LDS sample buffer.

Immunoprecipitates and whole-cell lysates were resolved by homemade SDS–PAGE gels or mPAGE® Bis-Tris precast gels (Millipore) and transferred onto nitrocellulose membranes using the Trans-Blot Turbo Transfer System (Bio-Rad). Membranes were blocked with 5% BSA in PBST (PBS containing 0.05% Tween 20) and incubated overnight at 4 °C with primary antibodies, followed by incubation with horseradish peroxidase (HRP)-conjugated secondary antibodies (Jackson ImmunoResearch) for 1 h at room temperature. Proteins were detected using enhanced chemiluminescence reagents (Thermo Fisher Scientific) or WESTAR NOVA 2.0 ECL (CYANAGEN).

### Cross-linking mass spectrometry

#### Sample preparation

Cross-linking reactions (100 μl) were performed in PBS with recombinant p100-C at a final concentration of 2 μM. Cross-linking reagent concentrations were optimized by titration to select conditions that avoided aggregation and minimized intermolecular cross-linking (Fig. S5D and E). Aliquots of the protein samples were treated with DHSO in combination with DMTMM at a 1:1 molar ratio, or with DMTMM alone, and incubated for 45 min at room temperature with shaking. DMTMM and DHSO were used at protein-to-cross-linker molar ratios ranging from 1:500 to 1:2,000. Reactions containing DHSO were quenched by adding Tris-HCl (pH 7.5), whereas reactions containing DMTMM alone were quenched by adding ammonium bicarbonate at an approximately 1:1 quencher-to-cross-linker molar ratio. All quenching reactions were incubated at room temperature for 30 min. A 10-μl aliquot of each sample was reserved for SDS-PAGE analysis.

For each condition, 6 μg of cross-linked material (DHSO or DMTMM linked samples) was processed using the modified S-trap method ^27^. Two digestion strategies were used for the DMTMM-crosslinked samples: digestion with trypsin alone and sequential digestion with trypsin and Asp-N. DHSO-crosslinked samples were digested sequentially with trypsin and Asp-N. Briefly, 10% SDS and 50 mM Tris-HCl (pH7.5) were added to each sample at 1:1 ratio. Samples were reduced with 5 mM dithiothreitol and alkylated with 10 mM iodoacetamide in the dark. Phosphoric acid was then added to a final concentration of 1.2%, followed by the addition of 90% methanol in 5 mM ammonium bicarbonate. Each sample was loaded onto an S-Trap 96-well plate (Protifi, USA) and digested overnight with trypsin (1:50 trypsin/protein) at 37 °C. The digested peptides were eluted using 50 mM ammonium bicarbonate; and Asp-N was added to this fraction and incubated for an additional 4 h at 37 °C. The trypsin-only DMTMM samples were not subjected to the additional Asp-N digestion. Two additional elutions were made using 0.2% formic acid and 0.2% formic acid in 50% acetonitrile. The three eluates were pooled and dried by vacuum centrifugation. Samples were kept at −20 °C until analysis.

#### Mass spectrometry analysis

UPLC/MS grade solvents were used for all chromatographic steps. Each sample was fractionated using splitless nano-ultra-performance liquid chromatography (15 kpsi M-Class nanoACQUITY; Waters, Milford, MA, USA). The mobile phases were A) H2O containing 0.1% formic acid and B) acetonitrile containing 0.1% formic acid. Samples were loaded into a Symmetry C18 reversed-phase trapping column (180 µm internal diameter, 20 mm length, 5 µm particle size; Waters). Peptides were subsequently separated using a T3 HSS nano-column (75 µm internal diameter, 250 mm length, 1.8 µm particle size; Waters) at a flow rate of 0.35 µL/min. Peptides were eluted into the mass spectrometer using the following gradient: 4% −30% B over 55 min, 30%-90% B over 10 min, held at 90% B for 7 min, and then returned to the initial conditions. Samples were run in a random order.

The M-Class nanoUPLC was coupled online via a nanoESI emitter (20 μm tip; Fossil Ion Tech, Madrid, Spain) to an Exploris 480 mass spectrometer (Thermo Scientific) equipped with a Nanospray Flex ion source (Thermo Scientific).

Data were acquired in data-dependent acquisition (DDA) mode using a Top-Speed method with a 2-s cycle time. MS1 spectra were acquired at a resolution of 120,000 (at m/z 200) over a mass range of 375-1500 m/z, with an AGC target of 200% and automatic maximum injection time. The minimum intensity threshold was set to 5 x 10^4^, precursor charges were limited to 3-8, and dynamic exclusion to 40s. MS2 spectra were acquired at a resolution of 30,000 with a quadrupole isolation width of 1.6 m/z, an AGC target of 75%, and a maximum injection time of 150 ms. HCD fragmentation was performed at 30 NCE, with the first mass set to 120m/z.

#### Data analysis

Raw mass spectrometry data were converted using MSConvert and searched against the p100 sequence using MeroX version 2.0.2.5. Precursor and fragment mass tolerances were set to 5 and 10 ppm, respectively. Searches were performed in quadratic mode with a minimum precursor charge state of 4+, peptide lengths of 4–30 residues, and up to four missed cleavages. Cleavage rules were specified after K and R and before D and E. Carbamidomethylation of cysteine was specified as a fixed modification, and oxidation of methionine as a variable modification. For DHSO, cross-linking sites were specified as D, E, and protein C termini on both sides. For DMTMM, K, S, T, Y, and protein N termini were allowed to pair with D, E, and protein C termini, using a zero-length cross-link definition corresponding to the loss of H O. Score cutoffs of 20 and 30 were applied to DHSO and DMTMM searches, respectively, with a minimum of two fragment ions per peptide. Target–decoy analysis was enabled with an FDR cutoff of 1%. Identified cross-links were subsequently inspected manually for spectral quality and consistency with the protease digestion used for each sample.

### Single-molecule FRET spectroscopy

#### Fluorescent labeling of proteins for smFRET

Purified proteins were first incubated with 2 mM DTT to reduce cysteine residues and subsequently desalted into labeling buffer (20 mM Tris, 50 mM KCl, pH 7.1). Labeling reactions were performed by first incubating protein samples with Alexa Fluor 488 maleimide (Invitrogen) at a 1:0.9 protein to dye molar ratio for 1 h at 25 °C, followed by incubation with Alexa Fluor 594 maleimide (Invitrogen) at a 1:1.3 protein to dye molar ratio for an additional hour at 25 °C. Finally, labeled proteins were separated from the unreacted dye using a HiTrap desalting column (GE Healthcare) and frozen in storage buffer (25 mM Tris, 150 mM NaCl, 5% (w/w) glycerol, pH 7.5). For fluorescence anisotropy experiments, single-dye labeled proteins were prepared by incubating the proteins with Alexa Fluor 488 maleimide at a 1:3 protein-to-dye molar ratio for 2 h at 25 °C.

#### Single-molecule FRET data acquisition and data analysis

Flow cells were prepared as described previously ^28^ and filled with a mixture of 30 pM labeled p100-C, 25 mM Tris-HCl (pH 7.5, Sigma Aldrich), 150 mM NaCl (J.T. Baker), 0.01% Tween (Thermo Fisher), either in the absence or presence of 1 µM NIK. As specified in Fig. S5H, the apparent K_D_ of the p100-C dimer is about 10-30 nM. At the concentration of 30 pM, <1% of p100-C is expected to be dimeric. An inverted confocal microscope ^29^ was used for data acquisition. Measurements were conducted in pulse-interleaved excitation mode ^30^ using a sequence of one pulse for donor excitation (485 nm, 30 µW) and one pulse for acceptor excitation (595 nm, 5 µW) at 20 MHz. The laser beams were focused 10 µm deep into the sample solution. As molecules diffused through the confocal volume, they generated short fluorescence bursts. Emitted photons were separated into donor and acceptor channels according to their wavelengths, using a dichroic mirror (T588lpxr; Chroma) and filtered by band-pass filters (ET 535/70m (Chroma) for the donor channel and ET 642/80m (Chroma) for the acceptor channel). Arrival times of these photons were registered by two single-photon avalanche photodiodes (SPCM-AQR-15; Perkin-Elmer) coupled to a time-correlated single-photon counting module (HydraHarp 400, PicoQuant).

To identify fluorescent bursts, the time interval between each pair of successive photons, Δ*t*, was calculated and smoothed using a moving-average window of 15 photons. Candidate bursts were defined as continuous photon streams with smoothed Δ*t* values below 20 μs. To be retained as a burst, each photon stream was additionally required to contain at least 30 consecutive photons with smoothed Δ*t* values below 10 μs. The raw FRET efficiency of each burst was calculated based on the photons detected in both channels following donor excitation only. The raw stoichiometry was obtained from the detected photons in both channels after both excitations, as described elsewhere ^31,32^. A 2D histogram of raw stoichiometry versus raw FRET efficiency was generated and used to determine donor leakage into the acceptor channel and direct excitation of the acceptor dye by the 485 nm-laser ^31^.

To isolate species containing a single donor and a single acceptor fluorophore, only photon bursts with the appropriate stoichiometry and fewer than 400 photons were selected. To exclude periods in which either fluorophore was non-emissive, the selected bursts were further refined by requiring the detection of donor photons at intervals of no more than 50 μs and acceptor photons at intervals of no more than 40 μs throughout each burst. Because acceptor-dark periods could result in artificially low FRET efficiency values, burst boundaries were trimmed to the first and last detected acceptor photons. The trimmed bursts were required to contain at least 30 photons detected after donor excitation. Following stoichiometry selection and burst refinement, typically 1000-5000 fluorescence bursts were retained from each experiment. Photon counts were subsequently corrected for background, donor leakage, and direct acceptor excitation, and a γ-factor of 0.9 was applied ^31,32^. Apparent distances between the labeled residues were calculated from the pooled corrected photon counts of all identified bursts using a Förster radius of 5.4 nm ^33^. These values represent apparent inter-dye distances and were not corrected for the length and conformational freedom of the dye linkers or other uncertainties associated with converting ensemble FRET efficiencies into absolute residue-to-residue distances. The shot-noise limited width was calculated to estimate how much of the width of the FRET efficiency histogram can be explained by the limited number of photons detected from each molecule, assuming all molecules have the same underlying FRET efficiency.

Four independent measurements were performed each in the absence and presence of 1 µM NIK. Increasing the NIK concentration did not further alter the FRET efficiency histograms, whereas higher NIK concentrations substantially increased the fluorescence background. To test whether the observed FRET efficiency changes could arise from nonspecific protein interactions or molecular crowding, we performed control measurements with 10 µM BSA, an unrelated protein present at a tenfold higher concentration than NIK. BSA did not reproduce the FRET efficiency changes induced by 1 µM NIK (Fig. S5B). Fluorescence anisotropy measurements further excluded changes in dye rotational freedom as the origin of the observed changes in the FRET efficiency histograms (see below).

We also considered the possibility that NIK could be oligomerized and promote intermolecular FRET by bringing two p100-C molecules into proximity. However, no increase in fluorescence burst brightness was observed upon NIK addition, arguing against NIK-induced association of multiple labeled p100-C molecules under the measurement conditions.

#### Steady-state fluorescence anisotropy

Steady-state fluorescence anisotropy measurements were performed on a Fluorolog-3 spectrofluorometer (Horiba) equipped with automated polarizers. Alexa Fluor 488-labeled protein samples (200 nM) were measured using a 5 s integration time. Samples were excited at 495 nm and the emission was recorded at 520 nm. Both the excitation and emission monochromators were set to slit widths of 5 nm. Anisotropy values were calculated using the built-in FluorEssence software. Three independently prepared samples were measured in the absence of NIK and three replicate measurements of the same sample were done in the presence of NIK. The alignment of the polarizers was verified by confirming that the anisotropy of light scattered by a dilute Ludox suspension (= 0.975 ± 0.002) was near unity. Measurements were performed on the single-cysteine variants C891, C703, and C615, as well as on the C432-C891 double-cysteine variant labeled only with Alexa Fluor 488. For C432, the double-cysteine variant was used rather than generating and purifying an additional C432 single-cysteine variant specifically for this control. Because the fluorescence anisotropy of the donor fluorophore at C891 was unaffected by NIK (Fig. S5C), the C432-C891 variant could still reveal that there are no NIK-dependent changes in dye rotational freedom at C432.

## Notes

### Competing Interest Statement

The authors have declared no competing interest.

