## Supplementary material for "In the non-canonical NF-κB pathway, binding of the initiating kinase NIK to the pathway’s target, p100, serves a nonenzymatic “licensing” role": Suppemental figures

### Figure S1

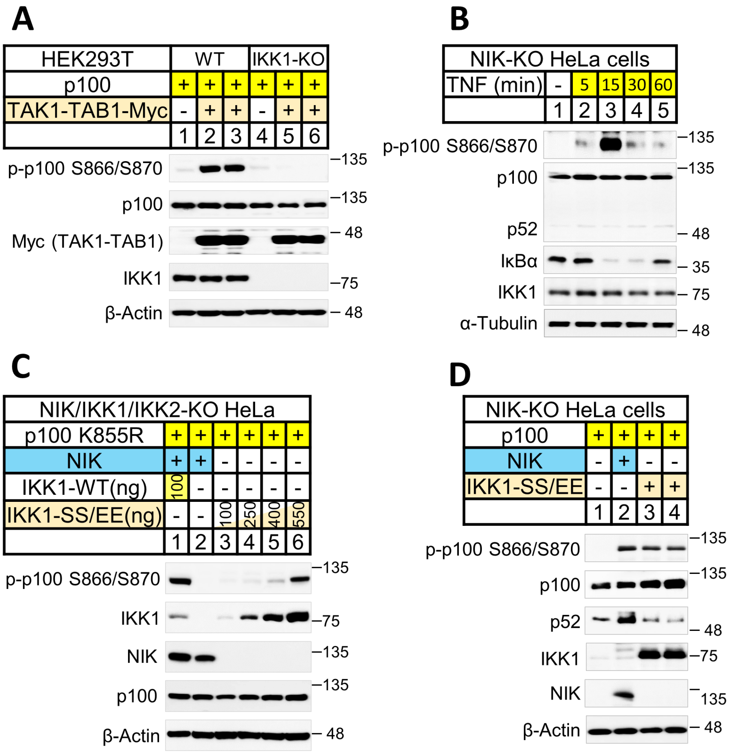
A
HEK293T
WT
IKK1-KO
p100
TAK1-TAB1-Myc
p-p100 S866/S870
p100
Myc (TAK1-TAB1)
IKK1
β-Actin
135
135
48
75
48
B
NIK-KO HeLa cells
TNF (min)
5
15
30
60
p-p100 S866/S870
p100
p52
IκBα
IKK1
α-Tubulin
135
135
48
35
75
48
C
NIK/IKK1/IKK2-KO HeLa
p100 K855R
NIK
IKK1-WT(ng)
IKK1-SS/EE(ng)
100
250
400
550
p-p100 S866/S870
IKK1
NIK
p100
β-Actin
135
75
135
135
48
D
NIK-KO HeLa cells
p100
NIK
IKK1-SS/EE
p-p100 S866/S870
p100
p52
IKK1
NIK
β-Actin
135
135
48
75
135
48

### Figure S2

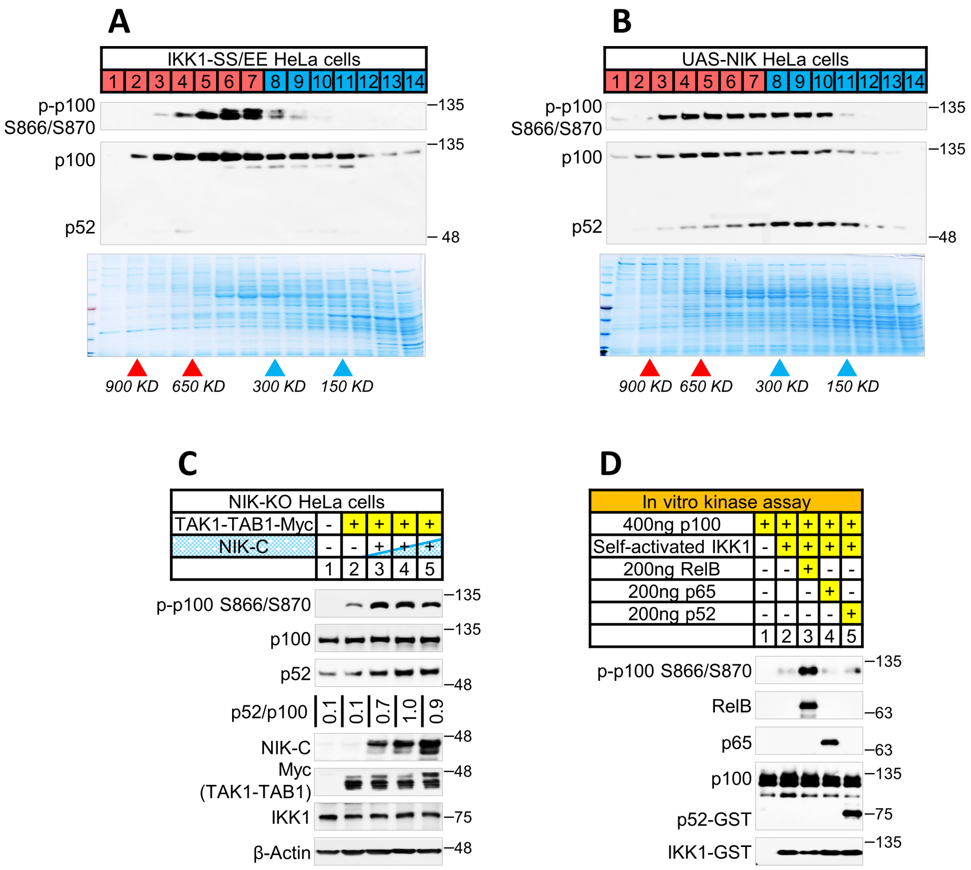
A
IKK1-SS/EE HeLa cells
1 2 3 4 5 6 7 8 9 10 11 12 13 14
p-p100 S866/S870
p100
p52
135
135
48
900 KD
650 KD
300 KD
150 KD
B
UAS-NIK HeLa cells
1 2 3 4 5 6 7 8 9 10 11 12 13 14
p-p100 S866/S870
p100
p52
135
135
48
900 KD
650 KD
300 KD
150 KD
C
NIK-KO HeLa cells
TAK1-TAB1-Myc - + + + +
NIK-C - - + + +
1 2 3 4 5
p-p100 S866/S870
p100
p52
p52/p100 0.1 0.1 0.7 1.0 0.9
NIK-C
Myc (TAK1-TAB1)
IKK1
β-Actin
135
135
48
48
48
75
48
D
In vitro kinase assay
400ng p100 + + + + +
Self-activated IKK1 - + + + +
200ng RelB - - + - -
200ng p65 - - + -
200ng p52 - - - - +
1 2 3 4 5
p-p100 S866/S870
RelB
p65
p100
p52-GST
IKK1-GST
135
63
63
135
75
135

### Figure S3

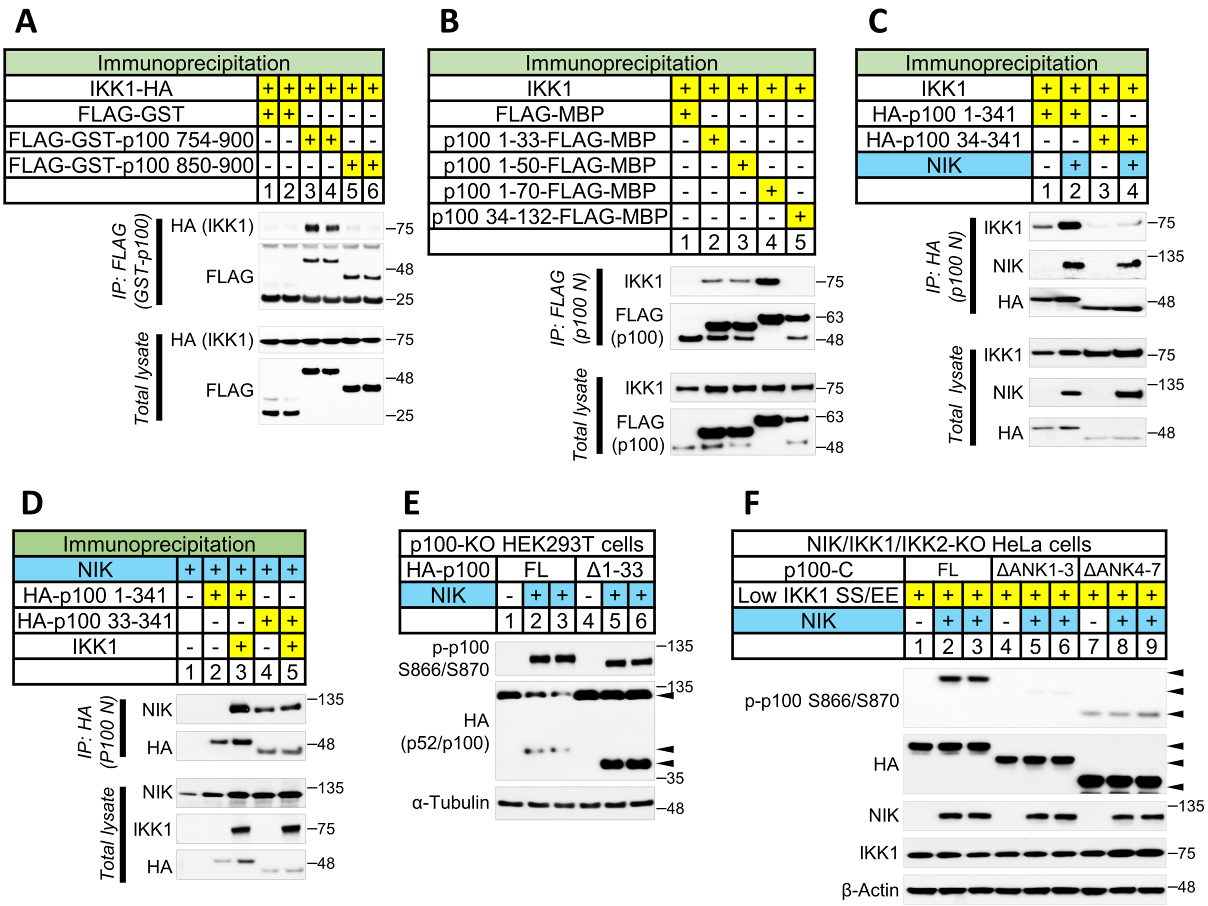

A
Immunoprecipitation
IKK1-HA
FLAG-GST
FLAG-GST-p100 754-900
FLAG-GST-p100 850-900
IP: FLAG (GST-p100)
HA (IKK1)
FLAG
Total lysate
B
IKK1
FLAG-MBP
p100 1-33-FLAG-MBP
p100 1-50-FLAG-MBP
p100 1-70-FLAG-MBP
p100 34-132-FLAG-MBP
IP: FLAG (p100 N)
FLAG (p100)
C
HA-p100 1-341
HA-p100 34-341
NIK
IP: HA (p100 N)
HA
D
HA-p100 33-341
IP: HA (P100 N)
E
p100-KO HEK293T cells
HA-p100
FL
Δ1-33
p-p100 S866/S870
HA (p52/p100)
α-Tubulin
F
NIK/IKK1/IKK2-KO HeLa cells
p100-C
ΔANK1-3
ΔANK4-7
Low IKK1 SS/EE
β-Actin

### Figure S4

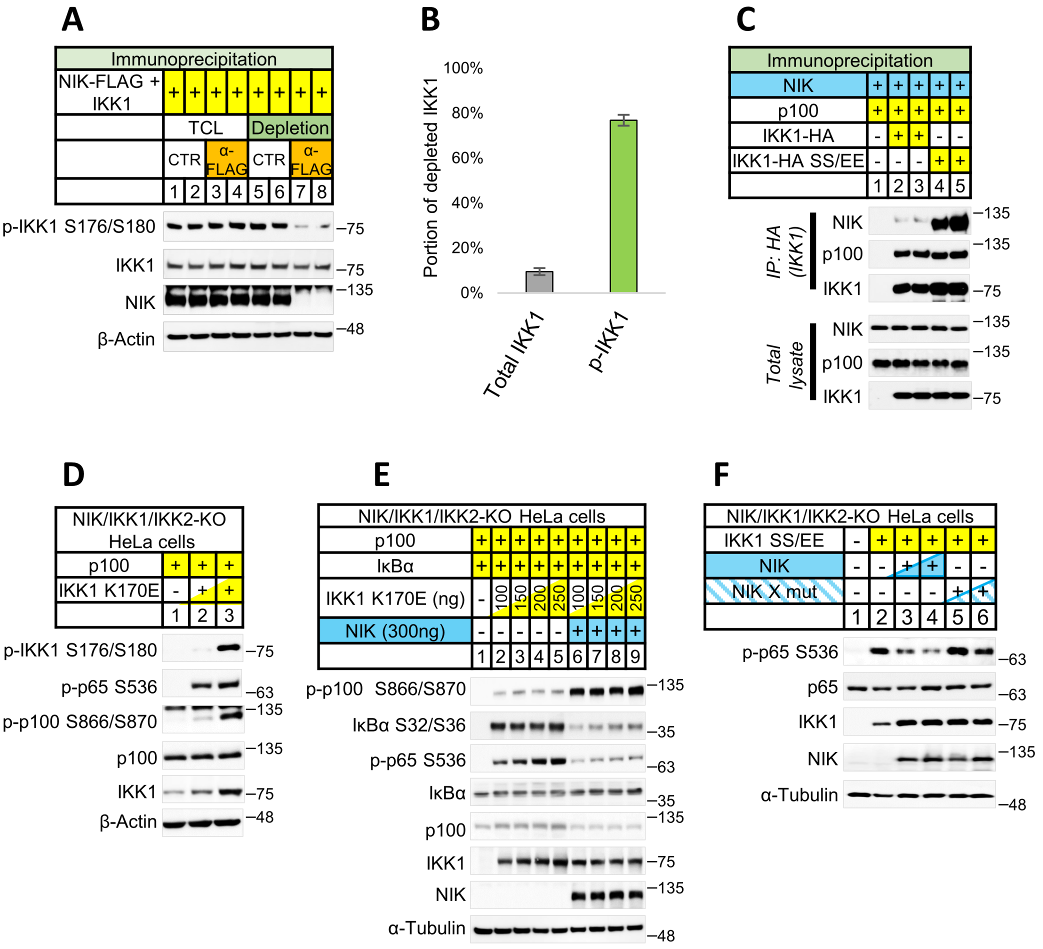
A
Immunoprecipitation
NIK-FLAG + IKK1
TCL
Depletion
CTR
α-FLAG
1 2 3 4 5 6 7 8
p-IKK1 S176/S180
IKK1
NIK
β-Actin
B
Portion of depleted IKK1
100%
80%
60%
40%
20%
0%
Total IKK1
p-IKK1
C
Immunoprecipitation
NIK
p100
IKK1-HA
IKK1-HA SS/EE
IP: HA (IKK1)
Total lysate
D
NIK/IKK1/IKK2-KO HeLa cells
p100
IKK1 K170E
p-IKK1 S176/S180
p-p65 S536
p-p100 S866/S870
E
IKBα
IKK1 K170E (ng)
NIK (300ng)
IkBα S32/S36
α-Tubulin
F
IKK1 SS/EE
NIK X mut
p65

### Figure S5

A

| In vitro binding assay | | | | | | |
|---|---|---|---|---|---|---|
| NIK | - | + | + | + | + | + |
| p100-C | + | + | - | - | - | - |
| p100-C C432-C891 | - | - | + | - | - | - |
| p100-C C615-C891 | - | - | - | + | - | - |
| p100-C C703-C891 | - | - | - | - | + | - |
| p100-C C738-C891 | - | - | - | - | - | + |
| | 1 | 2 | 3 | 4 | 5 | 6 |

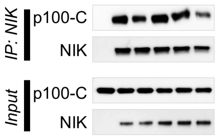


B

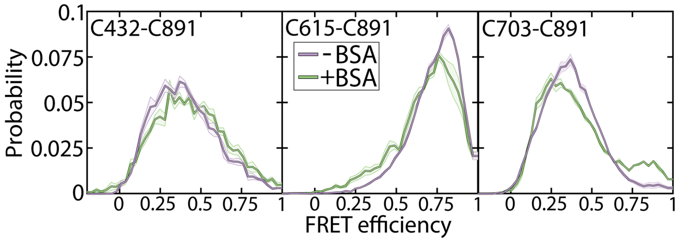


C

| Anisotropy | C432 | C615 | C703 | C891 |
|---|---|---|---|---|
| - NIK | 0.13±0.01 | 0.12±0.01 | 0.13±0.01 | 0.10±0.01 |
| + NIK | 0.13±0.01 | 0.12±0.01 | 0.12±0.01 | 0.09±0.01 |

D

| 2uM p100-C | | | | |
|---|---|---|---|---|
| DMTMM (mM) | - | 2 | 4 | 8 |
| DHSO (mM) | - | 2 | 4 | 8 |

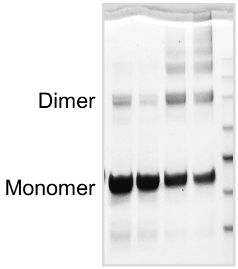


E

| 2uM p100-C | | | | | |
|---|---|---|---|---|---|
| DMTMM (mM) | 0.5 | 1 | 2 | 4 | 8 |

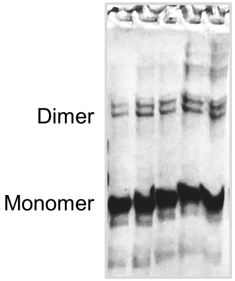


F

**Crosslinks of DeD with upstream reigons**

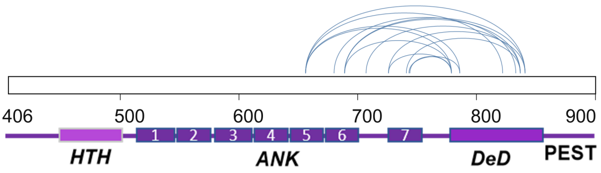


G

**Crosslinks of PEST with upsteam regions**

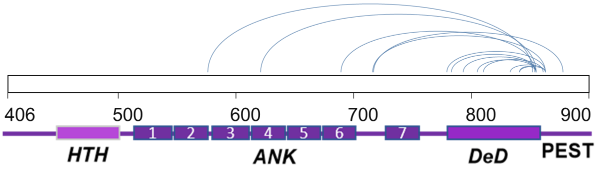


H

| p100-C | Monomer | Dimer | High-MW |
|---|---|---|---|
| 1nM | 82% | 15% | 3% |
| 10nM | 73% | 19% | 8% |
| 20nM | 69% | 24% | 7% |
| 30nM | 47% | 45% | 8% |
| Apparent $K_D$ | ~10-30 nM | | |
