## Supplemental table 2 for "In the non-canonical NF-κB pathway, binding of the initiating kinase NIK to the pathway’s target, p100, serves a nonenzymatic “licensing” role"

| **gRNAs** | **Seed sequences** |
| --- | --- |
| Human NIK-KO gRNA1 | TTTCAGAACGCATTTTCATC |
| Human NIK-KO gRNA2 | GAAAGCGTCGCAGCAAAGCC |
| Human NIK-KO gRNA3 | GCTCCTTCGGAGAGGTGCAC |
| Human IKK1-KO gRNA1 | GCTTGCCCTCCCGCCCCA |
| Human IKK1-KO gRNA2 | ACAGCGAGCAGATGACGTAT |
| Human IKK1-KO gRNA3 | GCCCCATGGAGCGGCCCCCG |
| Human IKK2-KO gRNA1 | AAATGCGGGAATCTTAACGC |
| Human IKK2-KO gRNA2 | GGGTGTCATAGCCCCGGGTT |
| Human IKK2-KO gRNA3 | TCCCGCATTTTAATGTTTTC |
| Human p100-KO gRNA1 | CAAATTGAACTCCTCCATTG |
| Human p100-KO gRNA2 | TACCAGGTCCACCTCGATCT |

**Table 2 gRNA seed sequences used in this study.**
